# The E2F4/p130 repressor complex cooperates with oncogenic ΔNp73α to promote cell survival in human papillomavirus 38 E6/E7-transformed keratinocytes and in cancer cells

**DOI:** 10.1101/2022.10.27.514150

**Authors:** Valerio Taverniti, Hanna Krynska, Assunta Venuti, Marie-Laure Straub, Cécilia Sirand, Eugenie Lohmann, Maria Carmen Romero-Medina, Stefano Moro, Alexis Robitaille, Luc Negroni, Denise Martinez-Zapien, Murielle Masson, Massimo Tommasino, Katia Zanier

## Abstract

Tumor suppressor p53 and its related proteins, p63 and p73, can be synthesized as multiple isoforms lacking part of the N- or C-terminal regions. Specifically, high expression of the ΔNp73α isoform is notoriously associated with various human malignancies characterized by poor prognosis. This isoform is also accumulated by oncogenic viruses such as Epstein–Barr virus (EBV), as well as genus beta human papillomaviruses (HPV) that appear to be involved in carcinogenesis. To gain additional insight into ΔNp73α mechanisms, we have performed proteomics analyses using human keratinocytes transformed by the E6 and E7 proteins of the beta-HPV type 38 virus as an experimental model (38HK). We find that ΔNp73α associates with the E2F4/p130 repressor complex through a direct interaction with E2F4. This interaction is favored by the N-terminal truncation of p73 characteristic of ΔNp73 isoforms. Moreover, it is independent of the C-terminal splicing status, suggesting that it could represent a general feature of ΔNp73 isoforms (α, β, γ, δ, ε, ζ, θ, η, and η1). We also show that the ΔNp73α- E2F4/p130 complex inhibits the expression of specific genes, including genes encoding for negative regulators of proliferation, both in 38HK and in HPV-negative cancer-derived cell lines. Consistently, silencing of E2F4 in 38HK and in cancer cells results in induction of senescence. In conclusion, we have identified and characterized a novel transcriptional regulatory complex that exerts pro-survival functions in transformed cells.

**IMPORTANCE:** The TP53 gene is mutated in about 50% of human cancers. In contrast, the TP63 and TP73 genes are rarely mutated but rather expressed as ΔNp63 and ΔNp73 isoforms in a wide range of malignancies, where they act as p53 antagonists. Accumulation of ΔNp63 and ΔNp73, which is associated with chemoresistance, can result from infection by oncogenic viruses such as EBV or HPV. Our study focuses on the highly carcinogenic ΔNp73α isoform and uses a viral model of cellular transformation. We unveil a physical interaction between ΔNp73α and the E2F4/p130 complex involved in cell cycle control, which rewires the E2F4/p130 transcriptional program. Consistently, we find that E2F4 gains pro-survival functions in transformed cells expressing ΔNp73α. This report shows, for the first time, that ΔNp73 isoforms acquire novel protein-protein interactions with respect to the TAp73 tumor suppressor. This situation is analogous to the gain-of-function interactions of p53 mutants supporting cellular proliferation.

## INTRODUCTION

The p53 family of proteins plays a key role in cancer prevention. It consists of three homologous transcription factors (TFs), namely p53, p63, and p73, which bind to common DNA promoter sites (p53RE). The TP53, TP63, and TP73 genes can be expressed as multiple isoforms lacking part of the N- or the C-terminal regions. In the case of TP73, the full-length TAp73 protein is transcribed from the P1 promoter upstream of exon 1, whereas ΔNp73 isoforms are generated from the P2 promoter within intron 3. These isoforms lack the N-terminal transactivation domain (TAD) and consequently act as dominant-negative inhibitors of full-length TAp73 and of its TAp53 and TAp63 homologues (1–3). Interestingly, the ΔN P2 promoter contains a p53 response element (RE) that enables TAp53 and TAp73 to induce expression of ΔN isoforms (4), thus creating a negative feedback loop that fine-tunes p53 and p73 functions. Additional isoforms of TAp73 and ΔNp73 proteins (i.e. α, β, γ, δ, ε, ζ, θ, η, and η1 isoforms) are generated by alternative splicing within the 3’ region of the gene (exons 10–14), giving rise to differences in the C-terminal regions of the proteins (5, 6).

Expression of the TA and ΔN proteins is highly cell- and tissue-specific. Whereas the ΔN isoforms support proliferation, the TA proteins promote cell-cycle arrest, senescence, and apoptosis, suggesting that the ratio between TA and ΔN proteins determines cell fate and oncogenesis (7). In healthy cells, where ΔNp73 levels are low, c-Abl phosphorylation coupled to Pin1 isomerase binding results in the stabilization of TAp73 (8–10), whereas ΔNp73 isoforms are rapidly degraded by other ubiquitin ligases that can discriminate between TA and ΔN isoforms (11) or *via* the calpain (12) and 20S proteasome (13) pathways.

Mutation of the TP53 gene is the most frequent genetic alteration, present in about 50% of human cancers. In contrast, the TP63 and TP73 genes are rarely mutated but are rather expressed as ΔNp63 and ΔNp73 isoforms in a wide range of human malignancies, which display unfavorable prognosis due to increased drug resistance (7, 14, 15). In particular, the ΔNp73α isoform, which comprises an intact (unspliced) C-terminal region, is upregulated in several cancers harboring wild-type TP53 and TP73 genes (including breast, prostate, liver, lung, and thyroid cancer), where it inhibits drug-induced apoptosis (15). Although the association of ΔNp73α with elevated carcinogenicity and chemoresistance can be explained by alterations of p53-regulated gene expression, it is not yet clear whether ΔNp73α could use additional mechanisms in promoting cellular transformation. Previous independent studies have shown that ΔNp73α is accumulated upon infection with the well-established oncogenic viruses, Epstein–Barr virus (EBV) (16) and other herpesviruses, such as the human cytomegalovirus (17). Accumulation of the ΔNp73α isoform is also induced in *in vitro* and *in vivo* experimental models by cutaneous beta (β) human papillomavirus (HPV) types (18–20), which appear to be involved in the development of skin squamous cell carcinoma (21).

In this study, we searched for protein factors that modulate the functions of ΔNp73α in transformed cells. We identified and characterized a novel interaction of ΔNp73α with the E2F4-5/p130 transcriptional repressor complex that modulates the cellular gene expression in *in vitro* HPV38 E6/E7-transformed human keratinocytes (38HK) as well as in cancer-derived cells.

## RESULTS

### ΔNp73α associates with the E2F4/p130 transcriptional repressor complex in 38HK

38HK are human keratinocytes immortalized by the E6 and E7 oncoproteins of cutaneous β-HPV38 (18). Here we used 38HK as a cellular model to perform proteomics analyses of ΔNp73α. The entire ΔNp73α sequence was fused to the N-terminus of the tandem affinity purification (TAP) tag, cloned in a lentiviral vector under the control of the weak promoter of Moloney murine leukaemia virus (22), and stably retrotransduced in 38HK (Fig. 1A). The nuclear fraction of 38HK expressing the ΔNp73α-TAP construct was recovered, and native ΔNp73α complexes purified using the TAP approach (23) (Fig. 1A and B). Finally, ΔNp73α binding partners were identified by mass spectrometry (S1 Table).

**Figure 1.**
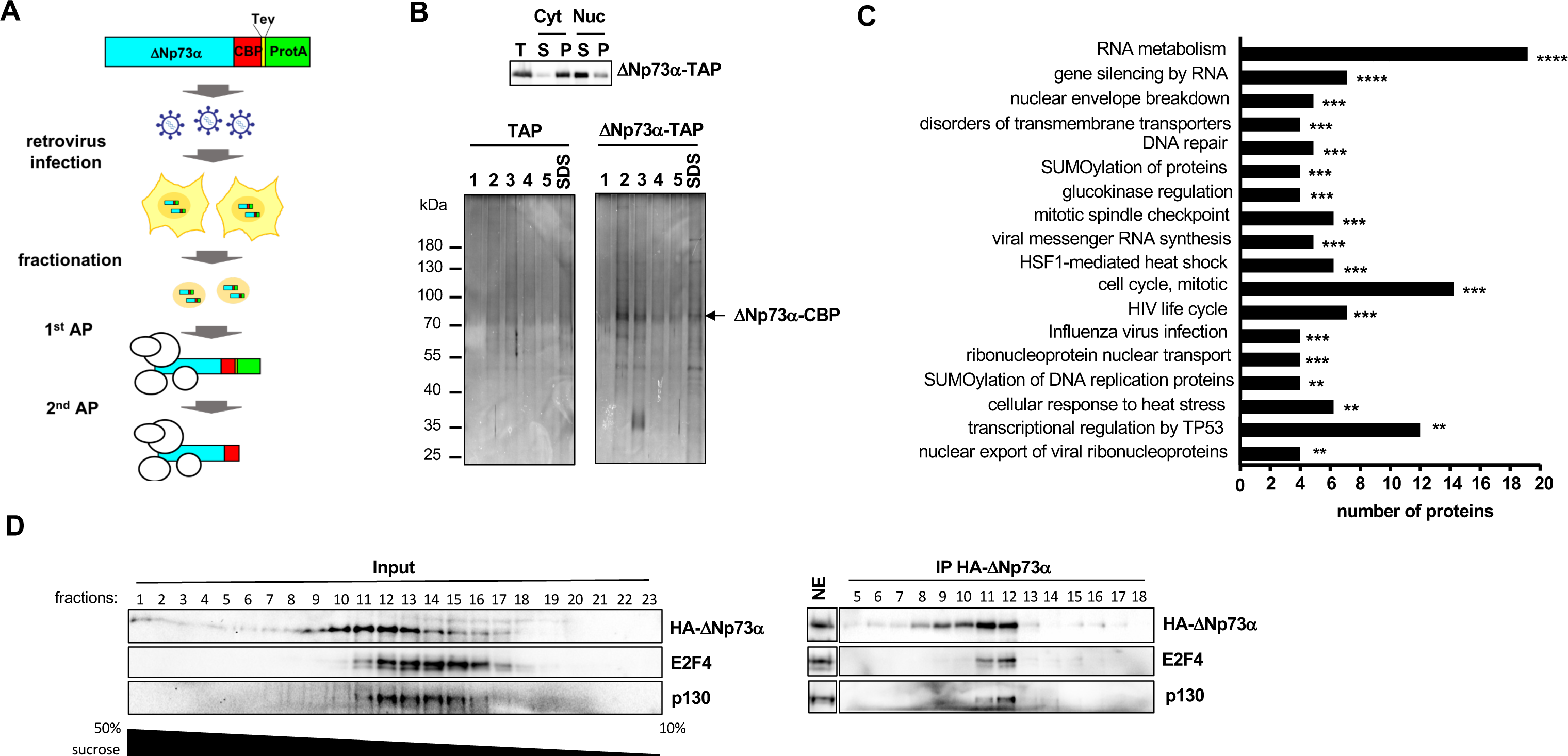
Proteomics analyses in 38HK identify the E2F4/p130 complex as a partner of ΔNp73α. **(A)** Illustration of the proteomics approach used to identify nuclear binding partners of ΔNp73α. (**B**) Expression and purification of ΔNp73α-TAP complexes. (*Upper panel*) Distribution of ΔNp73α-TAP in cytoplasmic and nuclear fractions of 38HK extracts. The two fractions were centrifuged and analyzed by Western blot using an anti-TAP antibody. T: total extract; S: supernatant; P: pellet. ΔNp73α-TAP is present mainly in the nuclear fraction (Cyt(P) and Nuc(S), see also Materials and Methods section). (*Lower panel*) Silver-stained 10% SDS-PAGE analysis of elution fractions 1–5 from the second affinity purification step (calmodulin resin) for ΔNp73α-TAP and control (TAP) purifications. An additional elution with SDS was performed to recover all the remaining proteins. (**C**) Pathway analysis of nuclear ΔNp73α binding partners using the Reactome database (73). Only significant pathways are shown (defined by a false discovery rate (FDR) value ≤ 0.02) and ranked based on the −log_10_ of the associated *P*-value (*: ≤ 0.05; **: ≤ 0.01; ***: ≤ 0.001; ****: ≤ 0.0001). Histogram bar size shows the number of input proteins involved in the corresponding pathway. See also S1 Table. (**D**) Sucrose gradient/co-IP experiments on endogenous 38HK proteins. (*Left panel*) Sucrose fractions of 38HK nuclear extracts stably expressing HA-ΔNp73α were migrated on a 10% SDS-PAGE gel and analyzed by Western blot using antibodies recognizing the HA tag, E2F4 and p130 proteins. (*Right panel*) Indicated fractions were immunoprecipitated using anti-HA conjugated beads. Unfractionated nuclear extract (NE) was loaded on the same gel as a reference for protein migration. Images for NE and sucrose fractions are derived from different exposures of the same membrane. See also original Western blot image on Mendeley data.

Bioinformatics analysis of the binders show an enrichment of proteins involved in transcriptional and cell-cycle control (Fig. 1C). In contrast, the E6 and E7 viral oncoproteins (38.E6 and 38.E7) do not appear to be associated with ΔNp73α.

Among the nuclear partners displaying high specificity for ΔNp73α-TAP, we identified the transcription factor E2F4 and the co-repressor retinoblastoma-like 2 (RBL2/p130) (S1 Table). To corroborate this finding, we performed co-immunoprecipitation (co-IP) experiments using nuclear extracts of 38HK stably expressing ΔNp73α fused to an HA tag (HA-ΔNp73α). Extracts were first applied to a 50–10% sucrose gradient and then incubated with anti-HA antibody conjugated beads. Results show that endogenous E2F4 and p130 coprecipitate with HA-ΔNp73α in fractions 11-12 containing all three proteins (Fig. 1D). By contrast, E2F4 and p130 are not recovered in fractions 13-16 presenting lower levels of HA-ΔNp73α, thereby confirming the co-IP specificity of factions 11-12. Consistently, incubation with an anti-E2F4 antibody coupled beads coprecipitates HA-ΔNp73α and p130 in both total and fractionated 38HK nuclear extracts (S1 Fig.). As an alternative approach, we also fractionated 38HK nuclear extracts by size exclusion chromatography, which provides more stringent complex fractionation conditions compared to sucrose gradients. Incubation with anti-HA beads coprecipitates endogenous E2F4 and p130 proteins in fractions containing the three proteins (S2 Fig.).

These results indicate that ΔNp73α interacts with several cellular proteins, including transcriptional regulatory complexes.

### ΔNp73α establishes direct protein–protein interactions (PPIs) with E2F4

E2F4 forms stable heterodimers with DP1/2 (24), whereas p73 hetero-tetramerizes with p63 (25, 26). Consistently, two unique DP1 peptides and several p63 peptides are detected in our proteomics analyses of ΔNp73α (S1 Table). To identify the direct binding partner of ΔNp73α, we evaluated binary interactions within a protein set that comprises ΔNp73α, TAp63α, E2F4, DP1, and p130 using the *Gaussia princeps* protein complementation assay (GPCA) (27). Here, the two test proteins are overexpressed in HEK293T cells as N-terminal fusions to the Gluc1 and Gluc2 inactive fragments of the *Gaussia princeps* luciferase. Based on previous benchmarking, protein pairs are considered as interacting if the normalized luminescence ratio (NLR) is above 5 (27). In our experiment, Gluc1-E2F4 and Gluc2-DP1 give a high binding response (NLR ∼ 500), which is consistent with the high affinity of the E2F4/DP1 heterodimer (Fig. 2A). As a control, we also measured the interaction between Gluc1-E2F4 and Gluc2-p130. The corresponding binding response (NLR > 40) indicates that, in the conditions of overexpression of this assay, only a fraction of Gluc2-p130 is sequestered by the competing E1A and SV40 Large T antigen proteins of HEK293T cells, leaving an excess of Gluc2-p130 free for interaction with E2F4. Besides these well-established interactions, E2F4 also binds to the ΔNp73α and the ΔNp73β isoforms (NLRs > 20) but not to p130 or DP1 (Fig. 2A). Moreover, no interaction is observed for TAp63α, ruling out any direct and sequence specific contribution of this protein in complex assembly.

**Figure 2.**
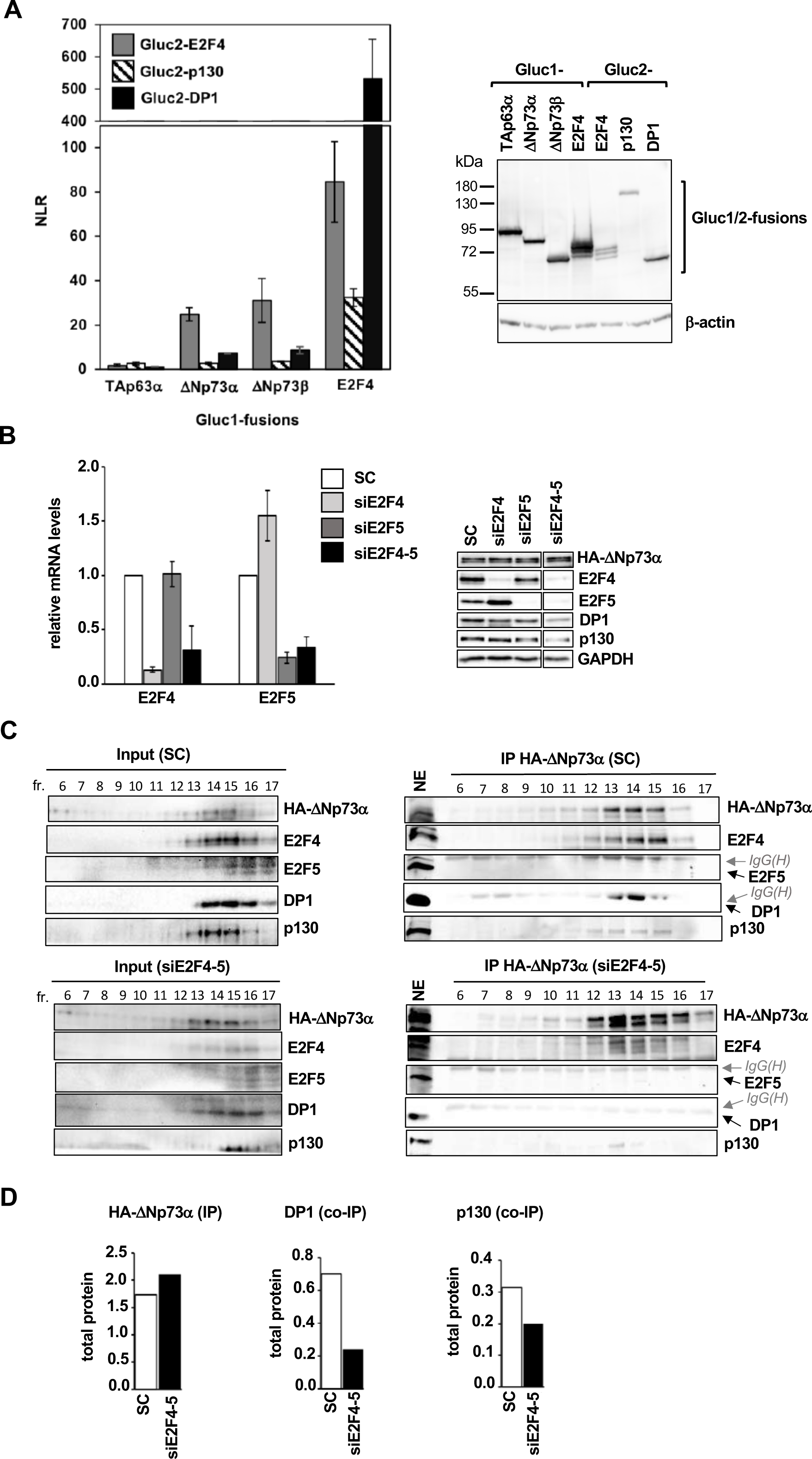
ΔNp73α establishes a direct PPI with E2F4. (**A**) (*Left panel*) Representative dataset for GPCA analysis of binary interactions between ΔNp73/TAp63 proteins and components of the E2F4/p130 complex. Pairwise combinations of Gluc1-ΔNp73/TAp63 and Gluc2- E2F4/DP1/p130 fusion constructs were co-transfected in HEK293T cells. After 48 hours, the luciferase activity was measured and expressed as normalized luminescence ratio (NLR) (27). The interactions of Gluc1-E2F4 with Gluc2-E2F4/DP1/p130 proteins are reported as internal controls. Error bars show standard deviations derived from triplicate measurements. (*Right panel*) 8% SDS-PAGE analysis of the expression of Gluc1-fused ΔNp73/TAp63 and E2F4 proteins, and of Gluc2-fused E2F4, DP1 and p130 proteins in HEK293T cells. Proteins were visualized by Western blot using an anti-Gluc antibody, which allows for detection of both Gluc1 and Gluc2 fragments, albeit with lower efficiency for Gluc2. The multiple bands detected for E2F4 likely correspond to multiple phosphorylation states of the protein as shown in (74). (**B**) Expression of DNp73a, E2F4-5, DP1 and p130 proteins in 38HK cells transfected with scramble (SC) siRNAs or siRNAs against either E2F4 (siE2F4) or E2F5 (siE2F5) or E2F4 plus E2F5 (siE2F4-5). (*Left panel*) mRNA levels measured by RT-qPCR. Error bars represent standard deviations derived from three independent experiments. (*Right panel*) Protein levels as determined by Western blot analysis. All samples were migrated on the same gel. See original Western blot image on Mendeley data. (**C**) Sucrose gradient/co-IP experiments using nuclear extracts from 38HK in the scramble (*upper panels*) and siE2F4-5 (*lower panels*) conditions. Indicated fractions were immunoprecipitated by anti-HA beads. HA-DNp73a, E2F4, E2F5, DP1, and p130 were detected by Western blot. IgG(H) chains migrate in close proximity with E2F5 and DP1 proteins. NE: unfractionated nuclear extract. See also S3 Fig. for experiments under single siE2F4 or siE2F5 conditions. See also legend of Fig. 1D. (D) Quantification of immunoprecipitated HA-DNp73a, DP1 and p130 proteins in the scramble and siE2F4-5 conditions of the co-IP experiments shown in panel C. For each protein, band intensities in the individual fractions are normalized to the intensity of the corresponding band in the NE control (Ifr/I_NE_). Then, the I_fr_/I_NE_ values of fractions 12 to 16 are summed to get an estimate of the total protein levels. The total DP1 and p130 protein values are additionally normalized to HA-DNp73a levels in the corresponding condition.

Next, we evaluated PPIs in conditions of protein partner depletion. Knockdown of E2F4 by small interfering RNAs (siRNAs) in 38HK leads to an increase in the functional homologue E2F5, and conversely, knockdown of E2F5 raises E2F4 levels (Fig. 2B). Immunoprecipitation of HA-ΔNp73α from fractionated nuclear extracts of 38HK treated with scramble siRNA or E2F4 siRNA leads to similar recoveries of p130 and DP1 (compare Fig. 2C *upper panels* with S3A Fig. *upper panels*). In contrast, simultaneous targeting of E2F4 and E2F5 by siRNAs strongly reduces E2F4 and E2F5 protein levels without affecting HA-ΔNp73α (Fig. 2B). In this condition, the ability of HA-ΔNp73α to coprecipitate DP1 and p130 clearly decreases (Fig. 2C *lower panels* and Fig. 2D).

Together, these results show that E2F4 establishes direct PPIs with ΔNp73α. In conditions of E2F4 knockdown, the E2F5 homologue is upregulated and interacts with ΔNp73α.

We then compared E2F4 and E2F5 for binding to ΔNp73α by GPCA in HEK293T cells. Based on the results obtained and taking into consideration the differences in the expression levels of the Gluc2-E2F5 and Gluc2-E2F4, we conclude that the two proteins are likely to interact with ΔNp73α with similar affinities under these assay conditions (S4 Fig.). However, endogenous E2F5 is not a partner of ΔNp73α neither in the proteomics analyses (S1 Table) nor in the co-IP experiments under the scramble condition (Fig. 2C *upper panels*). Hence, additional factors, such as expression levels or post-translational modifications (PTMs), may account for the preference towards endogenous E2F4 in 38HK cells.

### E2F4 discriminates between TAp73 and ΔNp73 isoforms

We reproducibly observed that the E2F4 binding response obtained with full-length TAp73α in GPCA experiments is lower (approximately 50%) than that obtained with the ΔNp73α isoform (Fig. 3A-B). This intriguing observation was confirmed by *in vitro* pulldown analyses that used recombinantly expressed full-length TAp73α and ΔNp73α proteins fused to the maltose binding protein (MBP) and clarified extracts of HEK293T cells overexpressing 3xFlag-E2F4 (Fig. 3C).

**Figure 3.**
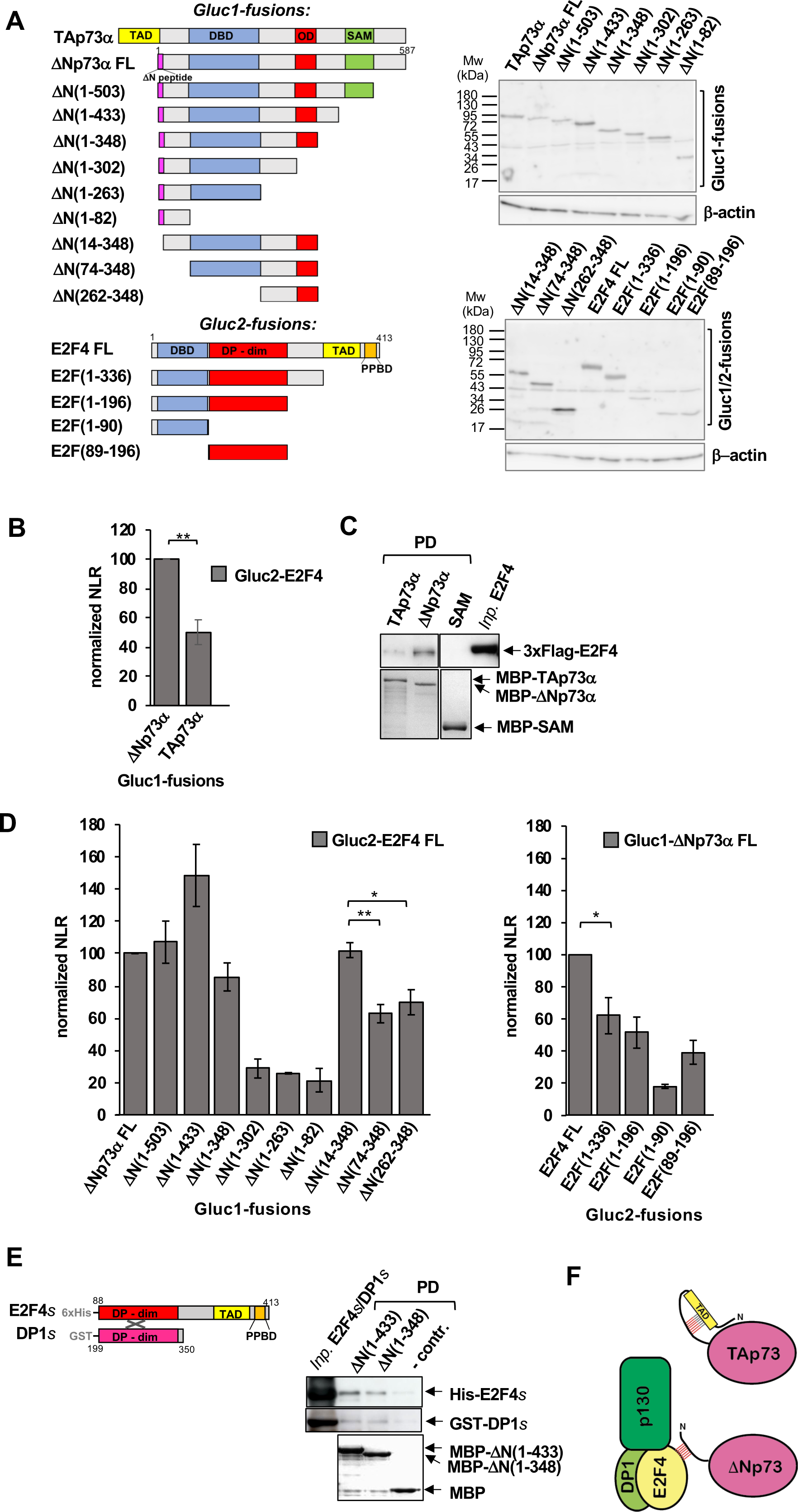
Identification of domains/regions mediating the ΔNp73α-E2F4 interaction. (**A**) (*Left panel*) Schematic representation of the Gluc1-fused TA/ΔNp73α and Gluc2-fused E2F4 constructs analyzed by GPCA. The ΔN(1–433) construct corresponds to the ΔNp73β isoform. Domains of TAp73α/ΔNp73α: transactivation domain (TAD, yellow); DNA-binding domain (DBD, blue); oligomerization domain (OD, red); sterile alpha motif domain (SAM, green); 13- amino-acid peptide specific for ΔNp73 isoforms (ΔN peptide, purple). Domains of E2F4: DNA-binding domain (DBD, blue); DP-binding and dimerization domain (DP-dim, red); transactivation domain (TAD, yellow); pocket protein binding domain (PPBD, orange). (*Right panel*) Expression levels of Gluc1- and Gluc2-fused TA/ΔNp73α and E2F4 constructs in HEK293T cells. Proteins were resolved on a 25%/10% gradient SDS-gel and visualized by Western blot using an anti-Gluc antibody. (**B**) GPCA analysis of the pairwise interactions of Gluc1-TAp73α or Gluc1-ΔNp73α FL with Gluc2-E2F4 FL. (**C**) MBP-pulldown analysis of the interactions of TAp73α and ΔNp73α FL with E2F4 FL. Amylose resin coupled to MBP- TAp73α or MBP-ΔNp73α FL constructs was incubated with clarified extracts from HEK293T cells transiently expressing 3xFlag-E2F4 FL. PD reactions were migrated on two separate 10% SDS-PAGE gels. One gel was used for E2F4 detection by Western blot using an anti-Flag antibody, the second gel for detection of MBP fusions by Coomassie staining. The negative control MBP-SAM construct (residues 405–502 of ΔNp73α) was migrated on the same gel as (see original Western blot images on Mendeley data). (**D**) GPCA analyses of pairwise ΔNp73α/E2F4 interactions. (*Left panel*) Gluc1-ΔNp73α deletion constructs versus Gluc2-E2F4 FL. (*Right panel*) Gluc1-ΔNp73α FL versus Gluc2-E2F4 deletion constructs. (**B** and **D**) The NLR values were normalized to the ΔNp73α FL/E2F4 FL interaction and are averages from three independent experiments (with each experiment being performed in triplicate). *P*- values are obtained from unpaired *t*-test, *n* = 3 biological triplicates (*: *P* < 0.05; **: *P* < 0.01). Interaction analyses using recombinant purified proteins. (*Left panel*) Schematic representation of the constructs used for the minimal E2F4*s*/DP1*s* heterodimer. (*Right panel*) Pulldown (PD) experiment using MBP-ΔN(1-433) (i.e. -ΔNp73β) and MBP-ΔN(1-348) or MBP (negative control) proteins and the minimal E2F4*s*/DP1*s* heterodimer. PD samples were migrated on two separate 10% SDS-PAGE gels. One gel was used for E2F4*s*/DP1*s* detection by Western blot using anti-His and anti GST antibodies, the second gel for detection of MBP-ΔNp73 fusions by Coomassie staining. (**F**) Proposed mechanism of ΔNp73α interaction with the E2F4/p130 complex. Red hatched box: E2F4 binding site.

To further characterize the interaction between E2F4 and ΔNp73α, we explored the binding properties of a panel of deletion constructs of ΔNp73α and E2F4 by GPCA (Fig. 3A). First, we searched for the region of ΔNp73α that interacts with E2F4. The C-terminal half of ΔNp73α (residues 349–587), comprising the sterile alpha motif (SAM) domain and the disordered linker and C-terminal regions, is dispensable for E2F4 binding, thus indicating that the interaction is independent of the C-terminal splicing status (α, β, γ, δ, ε, ζ, etc.) of ΔNp73 isoforms (Fig. 3D *left panel*, compare ΔNp73α FL with ΔN(1–348)). In contrast, deletion of the oligomerization domain (OD) strongly decreases binding, likely due to disruption of the tetrameric state and consequent loss of its avidity contributions (Fig. 3D *left panel*, compare ΔNp73α FL with ΔN(1–302)). Hence, the minimal ΔN(1–348) construct, which retains the binding properties of ΔNp73α FL, was further truncated from its N-terminus. Deletion of the first 13 residues specific to ΔNp73 isoforms does not affect the interaction. In contrast, deletion of the entire N-terminal disordered region (residues 1–73) reduces binding to 60%, that is to levels comparable to those of full-length TAp73α, whereas further deletion of the DNA-binding domain (DBD) does not have any effect (Fig. 3D *left panel*, compare TAp73, ΔNp73α FL with ΔN(14–348), ΔN(74– 348), and ΔN(262–348)). These results indicate that the N-terminal disordered region of ΔNp73 isoforms harbors a binding site for E2F4.

Then, we searched for the region of E2F4 that interacts with ΔNp73α. Deletion of residues 197–413 from the disordered C-terminus of E2F4, comprising the TAD and the pocket protein binding domain (PPBD), reduces binding activity to 60% (Fig. 3D *right panel*, compare E2F4 FL with E2F(1–336)), suggesting the existence of a binding site for ΔNp73 within this region. Further deletion of the C-terminus or of the DBD has only moderate effects on the interaction (Fig. 3D *right panel*, compare E2F4 FL with constructs E2F(1–196) and E2F(89– 196), which are probably related to the lower expression levels of these constructs (Fig. 3A *right panel*). In contrast, deletion of the DP-binding and dimerization (DP-dim) domain decreases binding to 20% (Fig. 3D *right panel*, compare E2F4 FL with E2F(1–90)). This domain of E2F4 provides avidity contributions but, unlike the OD of ΔNp73α, is also a well-known region for PPIs. Therefore, a contribution of the DP-dim domain towards ΔNp73 binding cannot be excluded.

To further corroborate these findings, we performed interaction analyses using recombinant purified proteins. Since E2F4 needs to be stabilized by the cognate DP1 partner in order to be purified, we used co-expression approaches in bacteria to produce a minimal E2F4/DP1 heterodimer (E2F4*s*/DP1*s*), which comprises the E2F4 regions required for the interaction with ΔNp73α (i.e. DP-dim and C-terminal regions, residues 88-413) and the DP-dim domain of DP1 (residues 199-350) (Fig. 3E *left panel*). In this way, we were able to obtain low concentration samples of E2F4*s*/DP1*s*, which were sufficient for pulldown analyses. Results from these analyses reproducibly show that E2F4*s*/DP1*s* interacts with the minimal MBP-fused ΔNp73 constructs (ΔN(1-433), equivalent to the ΔNp73β isoform, and ΔN(1-348)) but not with the MBP control (Fig. 3E *right panel*).

Taken together, our analyses suggest that the ΔNp73α-E2F4 interaction is mediated by a binding site within the N-terminal disordered region of ΔNp73 isoforms that is not accessible in TAp73 most likely due to intramolecular interactions (Fig. 3F).

### Expression and regulation of ΔNp73α-E2F4/p130 complex components in transformed *versus* untransformed cells

We evaluated the levels of ΔNp73α, E2F4 and p130 proteins in different primary keratinocyte (HPK) and transformed 38HK cell lines. Consistently with previous findings (18), ΔNp73α is observed in 38HK cell lines only, at both early (38HK(D2) and late (38HK(D3)) stages of transformation (Fig. 4A *upper panel*). In contrast, the E2F4 and p130 proteins are expressed in both primary HPK and 38HKs. λ- phosphatase treatment induces a small shift in the migration of p130, which suggests that this protein is phosphorylated to a certain degree in both HPKs and 38HKs (Fig. 4A *lower panel*). In line with these results, analysis of the cancer database shows that E2F4, and, albeit at a lower extent, the E2F5 homologue, as well as DP1 and p130 are similarly expressed in both normal and cancer tissues from different anatomical sites (S5 Fig.). Cellular fractionation experiments show that ΔNp73α, E2F4 and p130 mostly localize to the nucleus of 38HK and are equally distributed between the nuclear soluble (i.e. nuclear(S)) and chromatin fractions (Fig. 4B).

**Figure 4.**
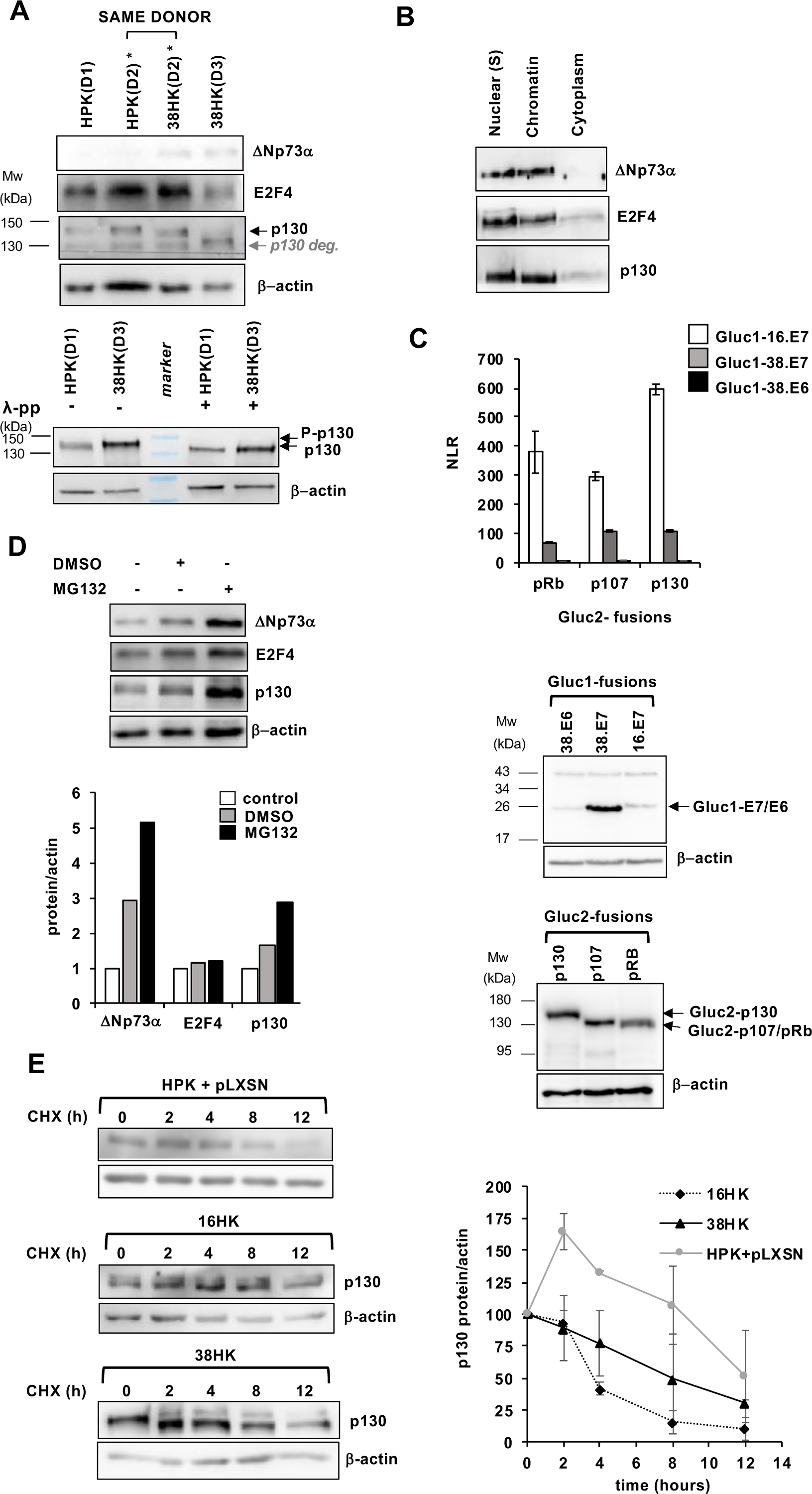
Expression of ΔNp73α-E2F4/p130 complex components in primary HPK and transformed 38HK cells. (**A**) (*Upper panel*) Immunoblotting of ΔNp73α and E2F4 and p130 proteins in human foreskin HPK and 38HK cell lines from 3 different donors (D1-D3). *: HPK(D2) and 38HK(D2) are from the same donor. 38HK(D2) is an early passage (i.e. passage 16) cell line, whereas the 38HK(D3) is a late passage cell line, which was used throughout this study. (*Lower panel*) Phosphorylation of p130. Total lysates of HPK and 38HK cells were incubated in presence or absence of λ-phosphatase (λ-pp) (8 U/μl). Note the small migration shift between the λ-pp– and λ-pp+ conditions. (**B**) Distribution of ΔNp73α and E2F4 and p130 proteins in the nuclear soluble (Nuclear(S)), chromatin and cytoplasm fractions of 38HK. (**C**) (*Upper panel*) Representative dataset of the GPCA analysis of the pairwise interactions between Gluc1-38.E7/E6 or Gluc1-16.E7 proteins and Gluc2-fused RB proteins co-expressed in HEK293T cells. Error bars show standard deviations derived from triplicate measurements. (*Middle and bottom panels*) Expression levels of Gluc1-38.E7/E6 and Gluc1-16.E7 proteins and Gluc2-RB proteins in HEK293T cells. Gluc1-38.E7/E6 and Gluc1-16.E7 were migrated on a 12% SDS-PAGE gel, whereas Gluc2-RB proteins on a 8% gel. (**D**) Comparison of ΔNp73α, E2F4 and p130 levels in 38(HK) under DMSO control *versus* MG132 treatment (4 hours incubation with 10 μM MG132) conditions. (*Left panel*) Western blot analysis. (*Right panel*) Quantification of Western blot bands showing protein levels normalized to actin. (**E**) Kinetics of p130 degradation in different keratinocyte cell lines. (*Left panel*) Western blot analysis of total lysates of 38HK, 16HK and primary HPKs stably transfected with pLXSN plasmid. All cells were treated with cycloheximide (10 μg/ml) during12 hours. Cells were collected at the indicated time points. (*Right panel*) p130 levels in the three cell lines at the different time points. The data reported are normalized to actin and to the values at time=0 (100%). Error bars represent standard deviation values derived from three independent experiments.

It is well-established that the HPV E7 oncoprotein binds to RB family members, inhibiting their interactions with E2F factors. Results from GPCA experiments show that, in spite of the high expression levels, 38.E7 binds to pRb, p107 and p130 with NLR values that are clearly lower compared to those displayed by E7 from the “high-risk” HPV16 virus (16.E7) with the same proteins (Fig. 4C). In addition to acting as a steric inhibitor, HPV E7 also promotes RB protein degradation *via* the proteasome. Treatment with the MG132 inhibitor induces a 2.5-fold increase in the levels of p130, whereas E2F4 levels remain unchanged (Fig. 4D). Analyses in conditions of cycloheximide treatment to block protein synthesis show that p130 degradation is significantly slower in 38HK cells as compared to keratinocytes transformed by the E6/E7 oncoproteins from HPV16 (i.e. 16HK, Fig. 4E). Together, these observations indicate that 38.E7 is less efficient in targeting p130 as compared to 16.E7, thus enabling the preservation of a nuclear pool of E2F4/p130, which becomes available for interactions with other factors. Finally, MG132 treatment of 38HK also leads to a 5-fold increase of ΔNp73α (Fig. 4D), an observation consistent with previous reports showing that ΔNp73α is subject to degradation in cells (11).

### E2F4-5 are required for survival of 38HK

We reasoned that the E2F4/p130 complex might acquire oncogenic functions by partnership with ΔNp73α, which is essential for survival of 38HK (18). Indeed, we observed that E2F4-5 knockdown by siRNAs in 38HK induces a different phenotype compared with the negative control (scramble siRNA), with cells appearing flattened and larger than normal 38HK. This phenotype is a feature of premature senescence. Consistently, assessment of β-galactosidase activity in 38HK shows that E2F4-5 knockdown leads to an increase in the number of senescent cells by a factor of two at 48 hours after transfection (Fig. 5A). To corroborate this finding, we performed immunofluorescent analysis of the H3K9Me3 (tri-methylation of Lys9 on histone H3) mark, which reports on the formation of senescence-associated heterochromatin foci (SAHF) (28). Single and double knockdown of E2F4 and/or E2F5 induces 2- and 4-fold increases in the percentage of 38HK cells positive for H3K9me3 marks, respectively (Fig. 5B). By contrast, E2F4-5 silencing in primary HPKs (S6 Fig.) does not affect SAHF formation (Fig. 5C). Furthermore, ectopic expression of E2F4 or E2F5 proteins, in cells where the E2F4 or of the E2F5 genes are silenced by shRNAs targeting the 3’UTR (S7 Fig.), rescues the proliferative phenotype of 38HK (S8 Fig.). This rules out the possibility that the senescence phenotype is an off-target effect of the knockdown approach.

**Figure 5.**
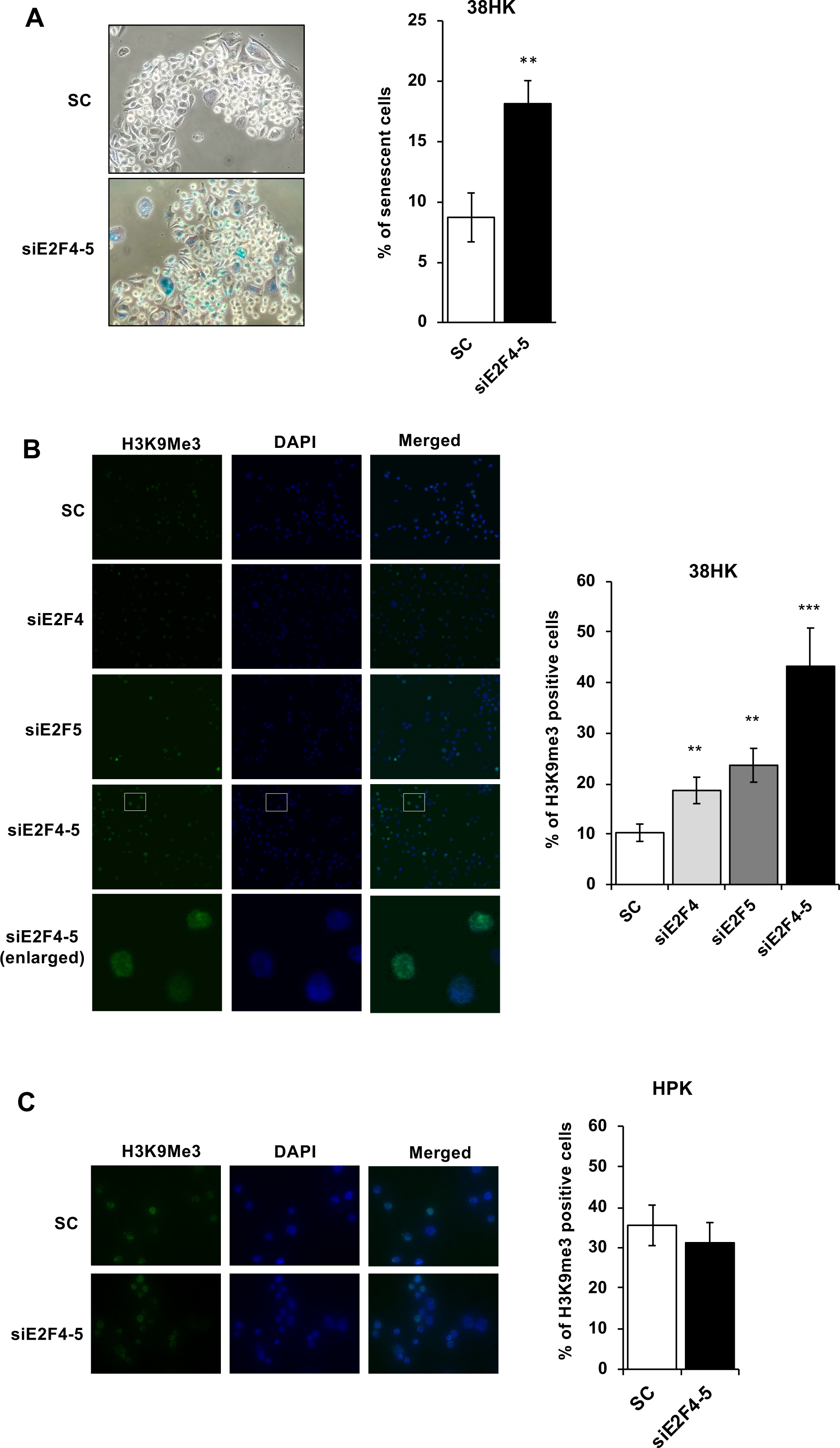
E2F4-5 depletion induces senescence in 38HK. (**A**) *In situ* senescence-associated β- galactosidase staining of 38HK. (*Left panel*) Representative photomicrographs of β- galactosidase staining at pH 6 on 38HK treated with SC or siE2F4-5 siRNAs at 48 hours after transfection. (*Right panel*) Percentage of senescent 38HK under scramble and siE2F4-5 knockdown conditions. (**B-C**) Immunofluorescent staining for SAHF in 38HK (**B**) and primary HPK cell lines (**C**). (*Left panels*)Representative images obtained using an epifluorescence microscope of 38HK and HPK cells transfected with SC, single siE2F4 or siE2F5, and double siE2F4-5 siRNAs. At 48 hours post-transfection, cells were layered on coverslips coated with polylysine and probed using an anti-H3K9me3 antibody followed by secondary Alexa Fluor 488-conjugated antibody. Nuclei were stained with DAPI (coloured blue). Images were merged using ImageJ software. Enlarged regions are indicated by white rectangles. (*Right panels*) Quantification of cells positive for H3K9me^3^ marks in the scramble and knockdown conditions performed using the ImageJ software. (**A**-**C**) The data are averages of three independent experiments with error bars representing standard deviation values. > 100 cells counted for each sample. *P*-values are obtained from unpaired *t*-test, *n* = 3 biological triplicates (**: *P* < 0.01; ***: *P* < 0.001). See also S6 Fig. for efficiency of E2F4-5 knockdown in HPKs.

These results point to pro-survival functions of E2F4-5 in transformed 38HK but not in primary HPKs.

### ΔNp73α cooperates with E2F4/p130 to inhibit gene expression

We used a three-step approach to search for genes that are targeted by both E2F4/p130 and ΔNp73α. First, we performed mRNA-seq analyses on 38HK in conditions of E2F4-5 depletion *versus* control. E2F4-5 knockdown by siRNAs reduces the levels of E2F4 and E2F5 proteins by 80% and 45%, respectively (S9A Fig.). This, in turn, leads to the upregulation of 2802 genes and to the downregulation of 2110 genes (adjusted *P*-value ≤ 0.05) (Fig. 6A, S2 and S3 Tables). Gene set enrichment analysis of upregulated genes identifies biological processes related to cell-cycle, tumor suppressor signaling, cancer, and viral infection pathways (Fig. 6B *left panel*). In contrast, downregulated genes are enriched in pathways linked to metabolism, DNA replication and extracellular matrix-receptor interactions (Fig. 6B *right panel*).

**Figure 6.**
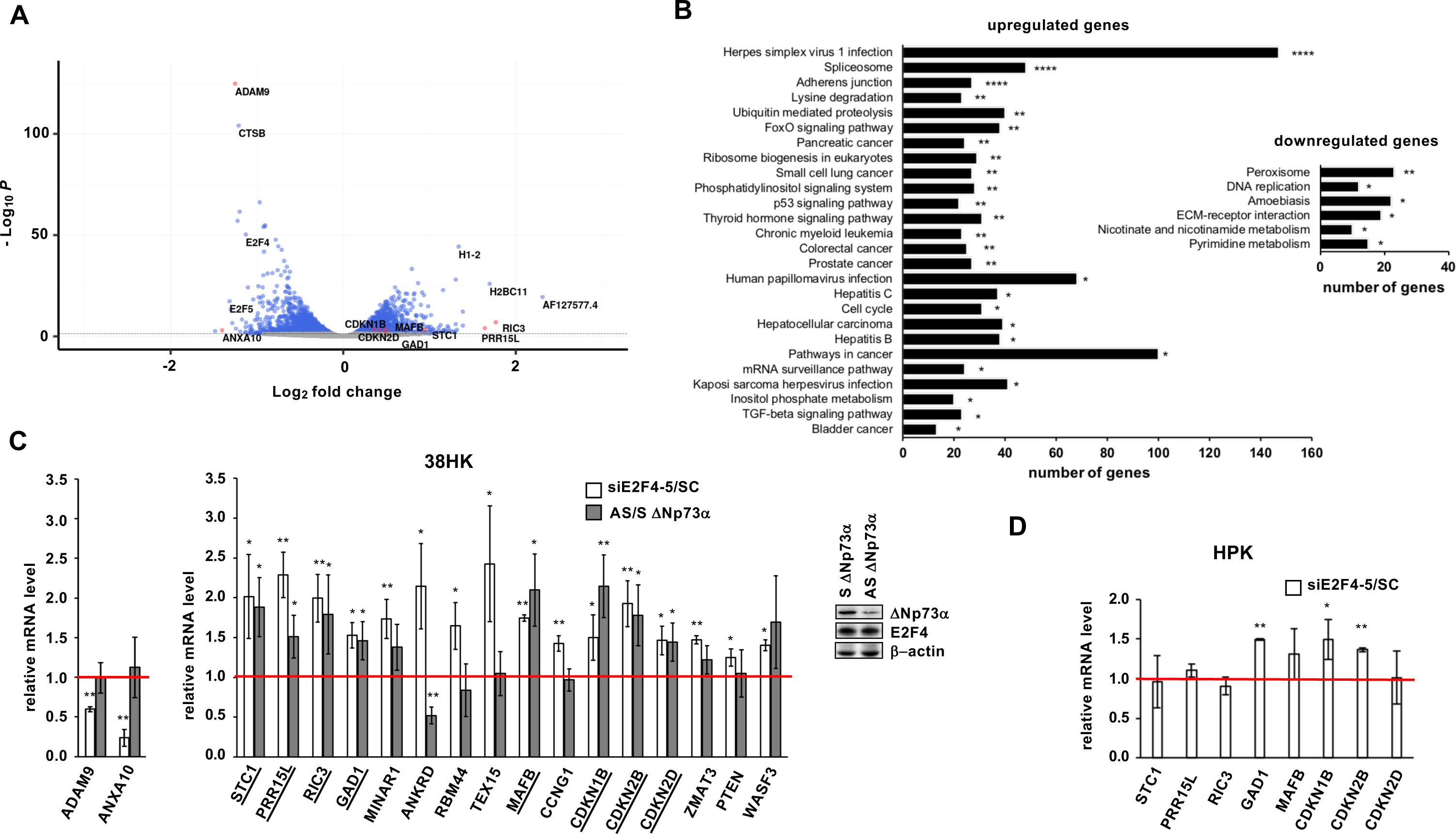
Identification of genes coregulated by E2F4-5 and ΔNp73α in 38HK. **(A)** Volcano plot of the genes deregulated by E2F4-5 knockdown in the mRNAseq analysis. *x*-axis: log_2_ transformation of the fold change; genes with a positive value are upregulated, whereas genes with a negative value are downregulated. *y*-axis: −log_10_ transformation of the adjusted *P*-value (*P*adj). Blue circles: significantly deregulated genes associated with *P*adj ≤ 0.05. Grey circles: non-significantly deregulated genes. Red circles: genes that are found to be coregulated by ΔNp73α and E2F4-5 and in panel C. **(B)** Biological processes enriched for the genes upregulated (*left panel*) and downregulated (*right panel*) in the RNA-seq analysis. Pathway analysis was performed using the KEGG-2019 database (75). Only significant pathways are shown (defined by *P*adj ≤ 0.05) and ranked based on the associated *P*-value (*: ≤ 0.05; **: ≤ 0.01; ***: ≤ 0.001; ****: ≤ 0.0001). Histogram bar size shows the number of genes involved in the corresponding pathway. (**C**) (*Left and middle panels*) Expression of selected genes in 38HK in conditions of E2F4-5 (white histograms) or ΔNp73α (grey histograms) depletion. mRNA levels are measured by RT-qPCR, first normalized to GAPDH expression and then to the SC or S control conditions. The red line refers to the mRNA level in the SC or S conditions (set to 1). “siE2F4-5/SC” and “AS/S ΔNp73α” values > 1 and < 1 indicate upregulation and downregulation of gene expression, respectively. Underlined genes are significantly upregulated upon either E2F4-5 or ΔNp73α knockdown (i.e. coregulated genes). (*Right panel*) ΔNp73α and E2F4 protein levels in 38HK in control and ΔNp73α knockdown conditions using S and AS oligos. See also S9B Fig. (**D**) Gene expression in primary HPKs in conditions of E2F4-5 knockdown. Only genes that are coregulated by E2F4-5 and ΔNp73α in 38HK (panel **C**) are investigated. (**C**-**D**) The values reported are averages of three independent experiments. *P*-values are obtained from unpaired *t*-test, *n* = 3 biological triplicates (*: *P* < 0.05; **: *P* < 0.01).

In the second step, we performed RT-qPCR validation of a sample of genes from the upregulated mRNA-seq dataset, which comprise or not p53/p73 response elements (REs) within their long promoter regions (defined as 2500 nucleotides upstream of the transcription start site [TSS]) (S4 and S5 Tables). A total of 16 genes were validated with significantly higher mRNA levels in the E2F4-5 knockdown condition compared to the scramble control (Fig. 6C *middle panel*, white histograms, siE2F4-5/SC > 1). In this validation we also included two genes from the downregulated mRNA-seq dataset (ADAM9 and ANXA10), for which we confirmed inhibition upon E2F4-5 depletion (Fig. 6C *left panel*, white histograms, siE2F4-5/SC < 1).

In the third step, we silenced ΔNp73α expression by treating 38HK with antisens (AS) oligonucleotides that target the N-terminal 13 aa peptide unique to ΔNp73 isoforms (Fig. 3A *left panel*). Notably, knockdown of ΔNp73α induces apoptosis in 38HK (18) and, for this reason, a compromise between cell viability and protein depletion needs to be reached. Here, we succeeded in decreasing ΔNp73α protein levels by about 50%, while leaving E2F4 levels unchanged (Fig. 6C *right panel* and S9B Fig.). ΔNp73α depletion does not affect the expression of ADAM9 and ANXA10 (Fig. 6C *left panel, gray histograms*). In contrast, it enhances the mRNA levels of 8 out of the 16 upregulated and validated genes (Fig. 6C *middle panel*, grey histograms, AS/S ΔNp73α > 1). These genes are STC1, PRR15L, RIC3, GAD1, MAFB, CDKN1B/p27^Kip1^, CDKN2B/p15^INK4b^ and CDKN2D/p19^INK4d^. Interestingly, three of these genes have no REs for p53/p73 within their long promoter regions (i.e. MAFB, GAD1, and CDKN2D/p19^INK4d^) (S5 Table). Together, these results provide evidence that E2F4/p130 and ΔNp73α coregulate a core set of genes in 38HK.

The changes in mRNA levels observed in these experiments are limited by the only partial depletion of E2F4-5 or ΔNp73α proteins that we were able to achieve. We also attempted different protocols for triple knockdown of ΔNp73α and E2F4-5 in 38HK, which all resulted in too high toxicity, thus preventing further gene expression analyses. To evaluate the synergy between these TFs, we analyzed the expression of the eight coregulated genes identified above albeit in primary HPKs, which do not express ΔNp73α. Results show that E2F4-5 silencing does not affect the expression of most of these genes (i.e. STC1, PRR15L, RIC3, MAFB, CDKN2D/p19^INK4d^) (Fig. 6D). This indicates that, in the absence of ΔNp73α, E2F4/p130 is unable to target these genes and to inhibit their expression.

To conclude, the synergy deriving from the ΔNp73α−E2F4 interaction results in the re- direction of the complex to genes that are different from those targeted by the individual components.

### The ΔNp73α-E2F4 interaction enhances recruitment to DNA regulatory regions

We evaluated the recruitment of the ΔNp73α-E2F4/p130 complex to the promoter regions of coregulated genes by chromatin immunoprecipitation (ChIP). For this, we selected STC1, whose promoter hosts both E2F and p53/p73 REs, and MAFB, which hosts E2F REs only (S5 Table). 38HK stably expressing HA-ΔNp73α were treated with scramble or E2F4-5 siRNAs, and processed for ChIP using anti-HA, anti-p73 and anti-E2F4 antibodies. The eluted DNA was analyzed by qPCR with primers flanking regions 1, 2, and 3 of the STC1 promoter (Fig. 7A), regions 1 and 2 of the MAFB promoter (Fig. 7B), or the negative control region reported in another transcriptomics study of E2F4 by Lee et al. (29) (Fig. 7C).

**Figure 7.**
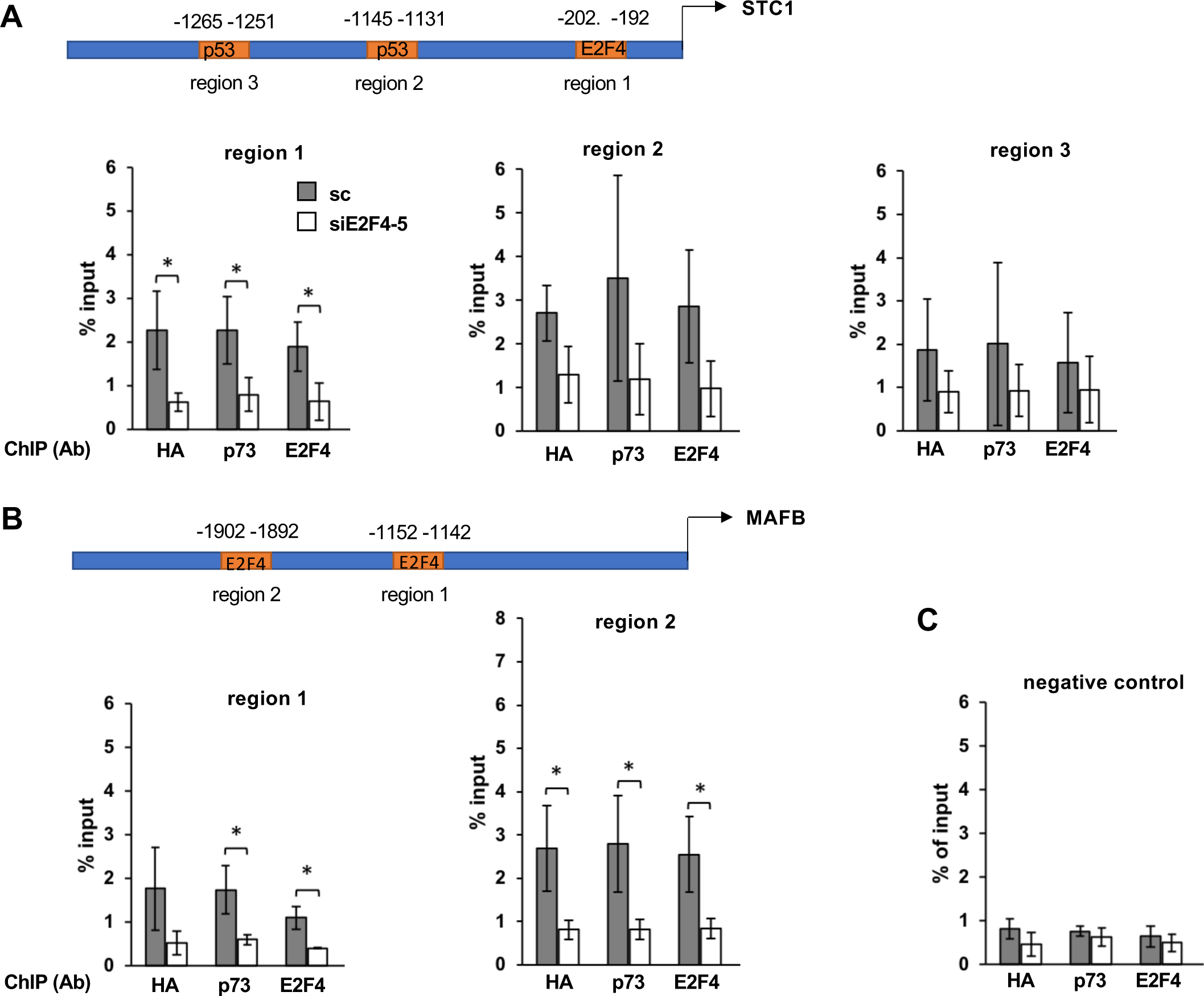
E2F4-5 knockdown decreases ΔNp73α recruitment to the promoters of the STC1 and MAFB genes. (**A, B**, *upper panels*) Schematic illustration of the long promoter regions (+2500 nucleotides from TSS) of the STC1 and MAFB genes. Numbers are the positions of TP53/TP73 and E2F4 REs associated with high prediction scores (see S4 Table). (**A, B,** *lower panels* and **C**) Results from ChIP experiments. 38HK expressing HA-ΔNp73α were treated with SC (grey histograms) or siE2F4-5 (white histograms) and harvested at 48 hours after transfection. Nuclear extracts were processed for ChIP using an anti-HA antibody recognizing HA-ΔNp73α (Ηα), an anti-p73 antibody recognizing both p73 and ΔNp73α proteins (p73) or anti- E2F4 (E2F4) antibodies. The eluted DNA was analyzed by qPCR with primers flanking regions 1, 2, and 3 of the STC1 promoter (**A**), regions 1 and 2 of the MAFB promoter (**B**), or the negative control region (S6 Table) (**C**). The amount of DNA bound by each protein is expressed as percentage of DNA in the input. (**A**-**C**) The data are averages of three independent experiments. *P*-values are obtained from unpaired *t*-test, *n* = 3 biological triplicates (*: *P* < 0.05).

Results show that HA-ΔNp73 and E2F4 are recruited to region 1 of the STC1 gene and to regions 1 and 2 of MAFB (Fig. 7, compare scramble conditions in panels A and B with scramble condition of panel C). Furthermore, E2F4-5 knockdown significantly reduces the recruitment not only of E2F4 but also of HA-ΔNp73α at these regions of the two genes.

These data indicate that ΔNp73α and E2F4 bind to the same promoter regions of coregulated genes. ΔNp73α recruitment to these regions takes place even in the absence of p53/p73 REs and is lost upon E2F4-5 knockdown, which demonstrates its dependence on the ΔNp73α-E2F4 interaction.

### Contributions of the ΔNp73α-E2F4/p130 complex in cancer cells

Because several studies have highlighted the importance of ΔNp73α in different types of cancer, we evaluated whether the ΔNp73α-E2F4 interaction occurs in cancer-derived cell lines. We previously identified an HPV-negative head and neck cancer cell line (HNC-136) and a breast cancer cell line (CAL- 51), both with high levels of ΔNp73α and wild-type TP53 (20). Nuclear extracts from HNC- 136 were treated by sucrose density gradient. Immunoprecipitation of endogenous ΔNp73α using anti-p73 antibody coupled beads leads to recovery of endogenous E2F4 and p130 proteins (Fig. 10A *lower panel*, fractions 15–17), thereby confirming ΔNp73α-E2F4/p130 complex formation in these cells. Interestingly, the complex from HNC-136 extracts migrates in fractions of lower sucrose density compared to the complex from 38HK extracts (compare Fig 10A with Fig. 1D). This suggests that ΔNp73α-E2F4/p130 may include auxiliary proteins that depend on the cellular context. In addition, although HNC-136 extracts exhibit clear expression of E2F5, no E2F5 is recovered upon ΔNp73α immunoprecipitation.

As in the case of 38HK, E2F4-5 depletion (S10A Fig.) leads to premature senescence in both HCN-136 and CAL-51 cell lines (Fig. 8B). Furthermore, E2F4-5 depletion in these cells increases the mRNA levels of most of the genes found to be coregulated by ΔNp73α and E2F4/p130 in 38HK (Fig. 8C, white histograms and S11 Fig.). In contrast, ΔNp73α depletion by antisens oligonucleotides induces massive cell death in both cancer cell lines. However, by carefully screening oligonucleotide concentration, we were able to identify one condition for the CAL-51 cell line, which limits the cell death (S10B Fig.) and results in a ΔNp73α knockdown of approximately 40% at 24 hours after transfection (S10C Fig.). Remarkably, even with such a partial depletion, we can observe an increase in the expression of several genes, including three genes (STC1, MAFB and CDKN2D/p19^INK4d^) that are also regulated by E2F4/p130 (Fig. 8C, grey histograms).

**Figure 8.**
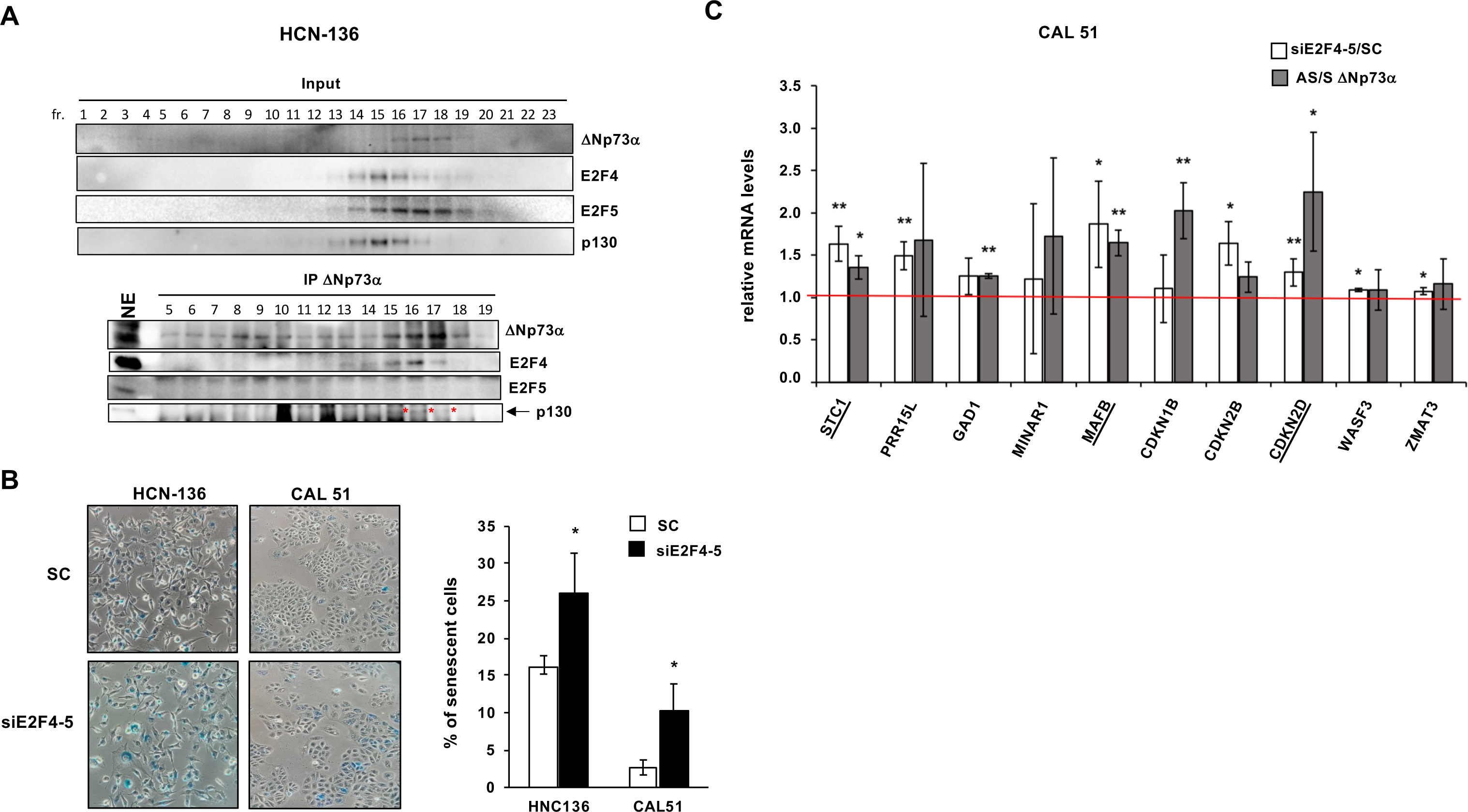
ΔNp73α cooperates with E2F4/p130 in HPV-negative cancer cells. (**A**) Sucrose gradient/co-IP experiments using nuclear extracts of HNC-136 cells. Sucrose fractions were immunoprecipitated using anti-p73 antibody-coupled beads and analyzed for endogenous ΔNp73α, E2F4, E2F5, and p130 proteins. Note that the p130 signal (highlighted by a red asterix) is partially masked by closely migrating non-specific band. See also expanded Western blot images on Mendeley data. See also legend of Fig. 1D. (**B**) *In situ* senescence-associated β- galactosidase staining of HNC-136 and CAL-51 cells (*Left panel*) Representative photomicrographs of β-galactosidase staining at pH 6 in cells treated with scramble or siE2F4- 5 at 48 h after transfection. (*Right panel*) Percentage of senescent cells under scramble and siE2F4-5 conditions. More than 100 cells were counted for each condition. (**B**) Expression profiles of selected genes in CAL-51 cells. mRNA levels determined by RT-qPCR under conditions of E2F4-5 (white histograms) or ΔNp73α (grey histograms) depletion. The red line refers to the mRNA level in the SC or S conditions (set to 1). “siE2F4-5/SC” and “AS/S ΔNp73α” values > 1 and < 1 indicate upregulation and downregulation of gene expression, respectively. The data are averages from three independent experiments, with error bars representing standard deviation values. *P*-values are obtained from unpaired *t*-test, *n* = 3 biological triplicates (*: *P* < 0.05; **: *P* < 0.01; ***: *P* < 0.001). See also Fig. S10 for E2F4-5 and ΔNp73α levels in the scramble and knockdown conditions.

Together, our data show that the ΔNp73α-E2F4/p130 complex regulates the expression of specific genes in non viral cancer cells.

### p19^INK4d^ expression induces senescence in 38HK

The CDKN2D/p19^INK4d^ gene is repressed by ΔNp73α and E2F4-5 in both 38HK and CAL51 cancer cell lines. A previous study showed that the p19^INK4d^ protein is involved in senescence by contributing to heterochromatin formation (30). To test whether similar mechanisms occur in our model system, we retro-transduced p19^INK4d^ or negative control pLXSN-GFP plasmid in 38HK (Fig. 9A) and performed senescence analyses. Results show that expression of p19^INK4d^ induces a 2-fold increase in the number of senescent cells in the β-galactosidase assay (Fig. 9B) and a 3-fold increase in the number of cells positive for the H3K9me3 marker of SAHF *foci* (Fig. 9C). This suggests that repression of the CDKN2D/p19^INK4d^ gene by the ΔNp73α-E2F4-5/p130 favors cell survival.

**Figure 9.**
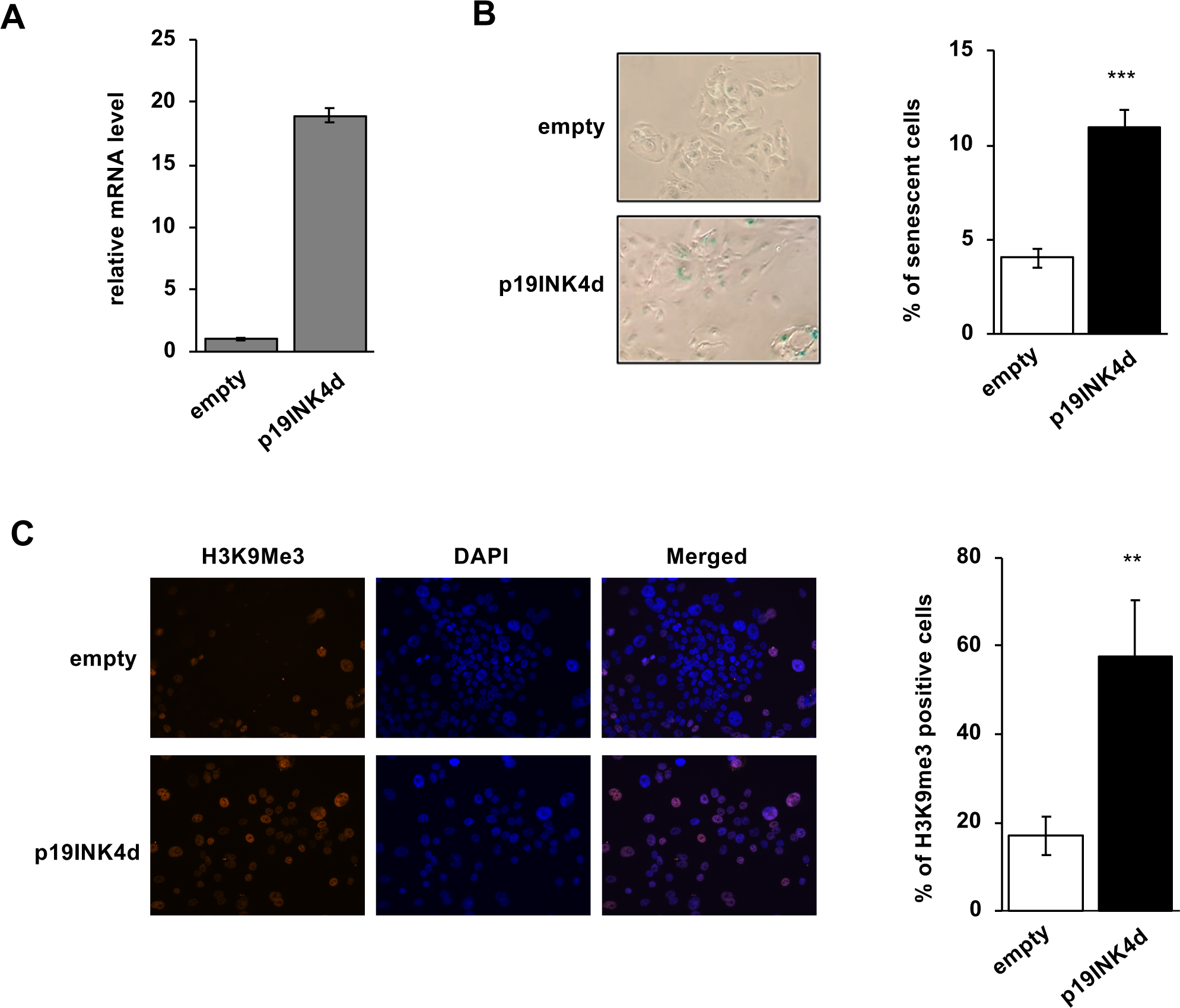
Expression of p19^INK4d^ induces senescence in 38HK. (**A**) p19^INK4d^ mRNA levels determined by RT-qPCR in 38HK retro-transduced with pMSCV-p19^INK4d^-IRES-GFP or with pLXSN-GFP (negative control). (**B**) *In situ* senescence-associated β-galactosidase staining. (*Left panel*) Representative photomicrographs of β-galactosidase staining at pH 6. (*Right panel*) Percentage of senescent 38HK in the negative control and p19^INK4d^ expression conditions. The data are averages and standard deviations from three independent experiments. (**C**) Immunofluorescent staining for SAHF. (*Left panel*) Representative immunofluorescence images. Cells were layered on coverslips coated with polylysine and probed using an anti- H3K9me3 antibody followed by secondary Alexa Fluor 555-conjugated antibody. Nuclei were stained with DAPI (coloured blue). (*Right panel*) Quantification of cells positive for H3K9me^3^ marks in the in the negative control and p19^INK4d^ expression conditions was performed using ImageJ software. (**A**-**C**) The data are averages from three independent experiments, with error bars representing standard deviation values. (**B-C**) More than 100 cells were counted for each condition. *P*-values are obtained from unpaired *t*-test, *n* = 3 biological triplicates (**: *P* < 0.01, ***: *P* < 0.001).

## DISCUSSION

In this study, we present what is, to the best of our knowledge, the first reported proteomics analysis on a ΔNp73 isoform, which has enabled us to identify the E2F4/p130 repressor complex as a nuclear partner of ΔNp73α. E2F4 belongs to the E2F family of proteins, whose members act as transcriptional activators (E2F1/2/3/6) and repressors (E2F4-5). E2F TFs associate with DP1–4 proteins through the dimerization domain and interact with the retinoblastoma (RB) family proteins (pRb, p130, p107) *via* the TAD. This interaction with RB proteins prevents recruitment of the transcriptional machinery. In the canonical model of cell-cycle progression, E2F4 and E2F5 mediate the repression of cell-cycle genes and cell-cycle arrest in the G0 phase and are part, together with p107/p130, DP1/2, and MuvB, of the DREAM complex (31, 32). Here, besides E2F4, DP1 and p130, none of the other DREAM complex subunits (i.e. LIN proteins, RBBP4) could be detected in association with ΔNp73α, suggesting that ΔNp73α-E2F4/p130 and DREAM are distinct complexes. In addition to the canonical functions, E2F4 has also activator functions that are RB independent (33). Here, E2F4-5 depletion in 38HK leads to downregulation of a large set of genes related to different biological pathways.

E2F4-5 and associated proteins (DP1 and p130) are expressed in both healthy and cancer tissue. In this work we unveil a direct PPI between E2F4 and ΔNp73α. Despite the homology with E2F5, E2F4 appears to be the preferred partner of ΔNp73α in transformed cells. This is in line with previous studies showing oncogenic functions for E2F4 but not for E2F5 (34–36).

Many of our analyses are performed in the 38HK model of viral transformation, in which ΔNp73α is the main expressed isoform, whereas TAp73 levels are under the detection limit (18). Neither the E6 nor the E7 viral proteins were detected among the ΔNp73α partners. Compared to the high-risk α-HPV16 E7 oncoprotein, β-HPV38 E7 appears to be less efficient in targeting p130, preserving, in this way, the integrity of a nuclear pool of E2F4/p130. Furthermore, the detection of a ΔNp73α-E2F4/p130 complex in extracts of the HPV-negative HNC-136 cancer cell line confirms that the ΔNp73α-E2F4 interaction is independent of the viral oncoproteins.

Results from binding analyses suggest that truncation of the TAD region of p73, giving rise to ΔN isoforms, leads to the exposure of a binding site for E2F4 and that this interaction mechanism is independent of the C-terminal splicing status of ΔNp73 isoforms. This finding strongly suggests that oncogenic ΔNp73 isoforms acquire novel PPIs with respect to TAp73, which are reminiscent of the gain-of-function PPIs established by oncogenic mutants of p53 with other TFs (37, 38). However, unlike mutant p53 interactions, which generally lead to activation of transcription, the interaction of ΔNp73α with the E2F4/p130 complex results in the inhibition of gene expression.

It is well established that direct PPIs between TFs belonging to different families enhance cooperative binding to DNA and that this can result in TF redirection to different sites (39–41). Here, we show that most genes coregulated by ΔNp73α and E2F4/p130 in 38HK are not targeted by E2F4/p130 in primary HPKs lacking ΔNp73α. In complementary ChIP experiments, we demonstrate that E2F4 mediates ΔNp73α recruitment to the promoter regions of two co-regulated genes. These findings provide evidence that the ΔNp73α-E2F4 interaction modulates the DNA binding specificity of both ΔNp73α and E2F4/p130.

Redirection arising from PPIs between TFs can have different physiological outcomes. In the case of most p53 gain-of-function interactions, this leads to the enhancement of cellular proliferation. Our finding on the senescence phenotype induced by E2F4-5 depletion points to pro-survival functions of E2F4-5 in transformed cells, which is in line with previous reports on the oncogenic functions of E2F4 in prostate (35) and breast (36) cancers. Interestingly, several of the genes coregulated by ΔNp73α and E2F4/p130 encode for senescence factors (i.e. CDKN1B/p27^Kip1^, CDKN2B/p15^INK4b^, CDKN2D/p19^INK4d^, STC1 and MAFB), which suggest that E2F4-5 pro-survival functions are related to partnership with ΔNp73α.

CDKN1B/p27^Kip1^, CDKN2B/p15^INK4b^ and CDKN2D/p19^INK4d^ belong to the *Cdkn* family. These genes encode for negative regulators of cell-cycle progression by binding and inhibiting cyclin-dependent kinases CDK1/2 (p27^Kip1^) and CDK4/6 (p27^Kip1^, p15^INK4b^ and p19^INK4d^), and represent well-established or candidate tumor suppressors (42–44). Multiple studies have shown that co-deletion of *Cdkn* genes in some human cancers are responsible for a worsen phenotype (45–47). p27^Kip1^ is regarded as an effector of senescence acting through the PTEN- p27^Kip1^ pathway (48, 49). Of note, CDKN1A/p21^cip1/warf1^, homologue of CDKN1B/27^Kip1^, is repressed by ΔNp73α but not by E2F4-5 in 38HK (*data not shown*). p15^INK4b^ plays a role in the maintenance of senescence and is transcribed from the INK4a/ARF/INK4b locus, which is deleted in a wide spectrum of tumors (50–53). p19^INK4d^ has also been shown to have antiproliferative properties and to be inactivation in human cancers (54–56). In agreement with a previous study (30), here we find that expression of p19^INK4d^ induces cellular senescence associated to heterochromatin formation in 38HK. In addition to *Cdkn* genes, STC1 and MAFB are also linked to senescence. Stanniocalcin 1, the product of the STC1 gene, is one of the most highly secreted factors in senescent cells (57). MAFB instead belongs to the AP1 family of TFs (58). Besides its well-established role as an oncogene, MAFB acts as a tumor suppressor in certain cellular contexts, by antagonizing oncoproteins such as HRAS and BRAF (59). Altogether our results on the repression of specific genes suggest that the interaction with oncogenic ΔNp73α alters the E2F4 biological properties in transformed cells. Yet, our data do not exclude that ΔNp73α or E2F4/p130 act alone or as part of other complexes also exerting oncogenic functions.

Our interaction analyses suggest that the E2F4 biding site is hindered in TAp73 by intramolecular interactions. p73 proteins are known to encompass a C-terminal transcription inhibitory domain (TID), which transiently interacts with the TAD domain (60). Deletion of the TID domain did not increase TAp73 binding to E2F4 (*data not shown*), suggesting that TID is not involved in the masking of the E2F4 interaction site. Future biochemical and structural studies are required to understand the precise mechanisms underlying the ΔNp73α-E2F4 interaction.

In conclusion, our study provides evidence that ΔNp73 isoforms acquire novel PPIs with other TFs. The ΔNp73α partner investigated here is E2F4, which gains oncogenic properties by supporting cell proliferation and survival.

## MATERIALS AND METHODS

### DNA constructs

The ORFs encoding *human* E2F4, *human* E2F5 and *human* DP1 were purchased from Addgene, whereas the ORF encoding human p130 from Origene.

#### Retroviral constructs

ΔNp73α-TAP was generated in two steps. First, the ΔNp73α open reading frame (ORF) was amplified by PCR and cloned in the pCDNA-TAP vector (23). Second, the pCDNA ΔNp73α-TAP plasmid was used as a template for PCR amplification of the ΔNp73α-TAP fusion, which was cloned into pBABE-puro retroviral vector (22) (pBABE ΔNp73α-TAP). For HA-ΔNp73α, the ΔNp73α ORF was amplified with an oligo complementary to its N-terminus and containing the HA tag and directly cloned into pBABE-puro (pBABE HA-ΔNp73α). pMSCV-p19Ink4d-IRES-GFP plasmid expressing *mouse* p19INK4d (sharing 87% sequence identity with human p19INK4d) (61) was obtained from Addgene.

#### Constructs for expression in mammalian cells

ORFs encoding for ΔNp73α, TAp73α, TAp63α, E2F4, E2F5, DP1, and p130 were amplified by PCR and cloned into the pDONR207 vector by recombination cloning (Gateway system, Invitrogen). The resulting pEntry clones were then transferred into the GPCA destination vectors pSPICA-N1 and pSPICA-N2 (27) and into a pcDNA3 vector enabling expression of an N-terminal 3xFlag tag.

#### Constructs for E.coli expression

ORFs encoding for ΔNp73α, TAp73α and constructs of these proteins were cloned in the NcoI and KpnI sites of a modified pETM-41 vector containing an N-terminal MBP tag followed by a TEV cleavage site. The E2F4(84–413) and DP1(199-350) DNA constructs for expression of the minimal E2F4*s*/DP1*s* heterodimer were cloned in the NdeI and BamHI sites of the pmCS and pnEA vectors, which allow for fusion to N-terminal 6xHis and GST tags, respectively (62).

#### shRNA constructs for gene knockdown

Oligonucleotides coding for shRNAs, which target the 3’-UTR of E2F4 and E2F5 genes (S6 Table), were cloned into the NdeI and EcoRI sites of the lentiviral expression vector pLKO.1. As a negative control a plasmid expressing the scramble sequence (MISSION pLKO.1-puro shRNA Control Plasmid DNA) was purchased from Sigma-Aldrich.

All constructs were verified by DNA sequencing.

#### Cell lines and cell culture

Phoenix, NIH 3T3, HEK293T, HNC-136 and CAL-51 cells were cultured in Dulbecco’s modified Eagle’s medium (DMEM), supplemented with 10% calf serum (NIH 3T3) at 37°C with 5% CO_2_.

38HK (63) were grown together with NIH 3T3 feeder layers in FAD medium containing 3 parts Ham’s F12, 1 part DMEM, 2,5% fetal calf serum, insulin (5 µg/ml), epidermal growth factor (10 ng/ml), cholera toxin (8.4 ng/ml), adenine (24 µg/ml), and hydrocortisone (0.4 µg/ml). Feeder layers were prepared by treating NIH 3T3 with mitomycin C for 2 hours. Around 3x10^5^ of treated NIH 3T3 were co-cultured with 38HK cells in T75 cell culture flasks. Feeder layers were removed by incubating the cell co-cultures with 5 ml of PBS 1X supplemented with 2mM EDTA. In this way, more than 95% of the feeder cells were removed. Then, 38HK cells were collected by scraping for analysis.

HPK cells were freshly isolated from neonatal foreskin and cultured in Keratinocyte Growth Medium 2 (PromoCell, Heidelberg, Germany).

#### Cell line generation

Retrovirus transduction system was used to generate 38HK cells stably expressing ΔNp73α- TAP, HA-ΔNp73α and p19INK4d constructs. High-titer retroviral supernatants were generated by transient transfection of Phoenix cells with the retroviral constructs described above and used to infect 38HK as described previously (64). Briefly, 500 μl of DNA mix (10 μg plasmid DNA, 248 mM CaCl_2_) were gently mixed to 500 μl of 2X HBS-buffered saline (1.5 mM Na2HPO4, 50 mM HEPES, 280 mM NaCl, 10 mM KCl, 12 mM Dextrose, pH 7.05). The transfection mix was then used to transfect Phoenix cells cultured in 5 ml fresh medium supplemented with 25 μM Chloroquine for 6/8 hours. After 48 hours, the culture medium containing the retrovirus was filtered (0.2 μm filter), mixed with 5 μl of polybrene (Sigma) and used to infect 38HK cell cultures for 3 hours. 24 hours after infection, 38HK were selected in 0.2 µg/ml of puromycin for 3-5 days.

#### Transfection conditions

38HK, HNC-136, and CAL-51 cell lines were transiently transfected with siRNAs and sense/antisense oligonucleotides (S6 Table) using Lipofectamine 2000 (Invitrogen). After 4 hours, the transfection mix was removed and the cells cultured in FAD medium (without antibiotic and cholera toxin). 38HK were transiently transfected with lentiviral shRNA plasmids using TransIT-Keratinocytes Transfection Reagent (Mirus) according to the manufacturer’s protocol. HEK293T cells were transfected using JetPEI® (Polyplus transfection).

#### Proteomics

38HK (about 2x10^8^ total cells) stably expressing ΔNp73α-TAP or TAP tag alone (negative control) were resuspended in 8 ml of cold buffer A (10 mM HEPES-KOH pH 7.9, 1.5 mM MgCl_2_, 10 mM KCl, 0.5mM DTT, 0.2 mM EDTA, protease inhibitor cocktail mix 1x, 10 mM NaF) and incubated for 30 minutes on ice. Then, cells were lysed by passing the mix through a 25 gauge needle 15 times and centrifuged for 10 minutes at 13400 rpm at 4°C. The supernatant (cytoplasmic soluble fraction) was flash-frozen and stored at -80°C, while the pellet (the nuclear fraction) was resuspended in 7 ml of cold buffer B (20 mM HEPES-KOH pH 7.9, 1.5 mM MgCl_2_, 250 mM NaCl, 20% glycerol, 0.5 mM DTT, 0.2 mM EDTA, protease inhibitor cocktail mix 1x, 10 mM NaF) and incubated on ice for 1 hour. The mix was centrifuged for 10 minutes at 13,400 rpm at 4°C and the supernatant (nuclear soluble fraction) was collected. Each lysis step was checked by Western blot (Fig. 1b, upper panel).

Buffer A was added to the nuclear soluble fraction to reach a final concentration of 200 mM NaCl and 16% glycerol. The resulting nuclear protein extract was transferred to an ultra-clear polycarbonate tube and centrifuged at 40,000 rpm for 1 hour at 4 °C using a SW41 rotor (Beckmann). After centrifugation, the supernatant was carefully collected and incubated overnight with 100 µl (dry bead volume) of prewashed IgG Sepharose beads. Beads were then washed 3 times with 10 ml of IPP150 buffer (10 mM Tris-Cl pH 8.0, 150 mM NaCl, 0.1% NP40) and once with 10 ml of TEV buffer (10 mM Tris-Cl pH 8.0, 150 mM NaCl, 0.1% NP40, 0.5 mM EDTA, 1 mM DTT). Subsequently, beads were resuspended in 1 ml of TEV cleavage buffer containing 15 µl (10 U/μl) of acTEV protease (Invitrogen) and incubated with gentle agitation for 4 hours at 16 °C. Protein complexes were eluted by gravity flow. Then, 3 ml of calmodulin binding buffer (10 mM β-mercaptoethanol, 10 mM Tris-Cl pH 8.0, 150 mM NaCl, 1 mM magnesium acetate, 1 mM imidazole, 2 mM CaCl_2_, 0.1% NP40) and 3 µl of 1 M CaCl_2_ were added to the 1 ml eluate containing the protein complexes to chelate the EDTA present in the TEV cleavage buffer. The resulting mix was incubated with 100 µl of prewashed (dry bead volume) calmodulin-Sepharose beads (Agilent Technologies) for 2 hours with gentle agitation at 4 °C. Beads were then washed 3 times with 10 ml of calmodulin binding buffer. The protein complexes were recovered with 5 consecutive elutions (200 µl each) with calmodulin elution buffer (10 mM β-mercaptoethanol, 10 mM Tris-Cl pH 8.0, 150 mM NaCl, 1 mM magnesium acetate, 1 mM imidazole, 2 mM EGTA, 0.1% NP40). An additional elution with 1% SDS was performed to recover all the remaining proteins (Fig. 1b, lower panel).

Elution fractions 2 from the ΔNp73α-TAP and TAP purifications were partially digested with trypsin and analyzed by LC-MS using an Orbitrap ELITE instrument equipped with a C18 Accucore 50 cm column. The generated data were analyzed using the Proteome Discoverer 2.4 tool. Proteins enriched more than 10-fold in ΔNp73α-TAP compared with the control (TAP-only) experiment are listed in S1 Table. Enrichment is calculated from the ratio of the sums of peptide peak areas in test and control experiments.

#### Cellular fractionation

38HK (out 2.5x10^7^ total cells) stably expressing HA-ΔNp73α were resuspended in 1 ml of cold buffer A (described in the proteomics section) and incubated for 15 minutes on ice. Then, cells were lysed by passing the mix through a 25 gauge needle 15 times and centrifuged for 5 minutes at 12000 rpm at 4°C. The supernatant, corresponding to the cytoplasmic soluble fraction, was recovered, while the pellet (the nuclear fraction) was resuspended in 200 μl of cold buffer B (described in the proteomics section) and incubated on ice for 30 min. Then, the mix was passed through a 27 gauge needle 15 times and centrifuged for 10 minutes at 12000 rpm at 4°C. The supernatant (i.e. the nuclear extract) was recovered and used for the co-IP experiments.

#### Sucrose gradient/co-immunoprecipitation (co-IP)

Sucrose density gradients were performed as previously described (65) with minor modifications. Briefly, step gradients were made by superposing sucrose solutions of different concentration (50%, 40%, 30%, 20%, 10%) in an ultra-clear polycarbonate tube (Beckman), and a linear gradient was allowed to form overnight at 4 °C. Then, 1.5-2 mg of nuclear extracts from 38HK HA-ΔNp73α or HNC-136 cells were carefully transferred to the top of the sucrose gradient and the protein complexes were separated based on their molecular weight by ultracentrifugation at 35,300 rpm for 16 hours at 4°C using the SW41 rotor. After centrifugation, 500 µl fractions were collected from the bottom of the tube by gravity flow.

ΔNp73α complexes were immunoprecipitated from each fraction using 30 μl of slurry anti-HA-agarose beads (Sigma-Aldrich, ref. A2095) or 20 μl of pre-coupled anti-E2F4 or anti-p73 Sepharose beads (dry bead volume). Briefly, each fraction was incubated with the beads for 2 hours (HA-beads) or 4 hours (E2F4 and p73-beads). After incubation, the beads were washed 5 times with 1ml of washing buffer (20mM Tris-HCl pH 7.5, 1.5 mM MgCl_2_, 150 mM NaCl, 0.2 mM EDTA, 0.1% Igepal). The protein complexes were eluted in 1x loading dye buffer.

#### GPCA assay

HEK293T cells were transfected using the reverse transfection method. Transfection mixes containing 100 ng of pSPICA-N2 and 100 ng of pSPICA-N1 plasmids expressing test proteins plus JetPEI® (Polyplus transfection) were dispensed in white 96-well plates. HEK293T cells were then seeded on the DNA mixes at a concentration of 4.2 x10^4^ cells per well. At 48 hours after transfection, cells were washed with 50 µl of PBS and lysed with 40 µl of *Renilla* lysis buffer (Promega, E2820) for 30 minutes with agitation. *Gaussia princeps* luciferase enzymatic activity was measured using a Berthold Centro LB960 luminometer by injecting 50 µl per well of luciferase substrate reagent (Promega, E2820) and counting luminescence for 10 seconds. Results are expressed as a fold change normalized over the sum of controls, specified herein as normalized luminescence ratio (NLR) (27). For a given protein pair A/B, NLR = (Gluc1-A + Gluc2-B) / [(Gluc1-A + Gluc2) +(Gluc1 + Gluc2-B)].

#### b-galactosidase and SAHF (senescence-associated heterochromatin foci) staining for senescence analyses

38HK, HNC-136, and CAL-51 cells were transiently transfected with siRNAs or lentiviral shRNA plasmids and cultured for 48 hours. 38HK transduced with p19INK4d were instead cultured for 72 h. The NIH 3T3 feeder layer was removed with PBS/EDTA from 38HK cultures prior to senescence analyses.

Senescence was assessed using the Senescence β-Galactosidase Staining Kit at pH 6 following the manufacturer’s instructions (Cell Signaling Technology). For SAHF staining, 38HK cells were layered on slides coated with polylysine and fixed in 4% paraformaldehyde in PBS (pH 7.4) for 15 min at room temperature, and permeabilized with 0.1% Triton X-100 in PBS for 15 min (66). Cells were incubated with H3K9me3 antibody (abcam; ab1220) for 2 hours at room temperature, followed by incubation with Alexa Fluor 488-conjugated or Alexa Fluor 555- conjugated secondary antibody for 1 hour at room temperature and mounted using Vectashield Antifade Mounting Medium with DAPI. The slides were visualized using a Nikon Eclipse Ti wide-field inverted fluorescence video microscope. The images thus captured were analyzed by NIS-Element software from Nikon.

#### Protein expression in *E. coli* and pulldown assays

The minimal E2F4*s*/DP1*s* heterodimer was produced by co-expression of 6xHis-E2F4 (84-413) and GST-DP1(199-350) in *E. coli* BL21 DE3 cells overnight at 15 °C. The bacterial pellet (500 ml expression) was resuspended in lysis buffer (20 mM Tris pH 8.0, 400 mM NaCl, 10% glycerol, 5mM DTT, lysozyme, 100 μg/ml DNAse I, 100 μg/ml RNAse, cOmplete EDTA-free (Roche)) and lysed by sonication. Cleared extracts were applied to Ni^2+-^NTA resin previously equilibrated in buffer A (20 mM Tris pH 8.0, 400 mM NaCl, 10% glycerol, 2mM DTT). After extensive washing the E2F4/DP1 heterodimer was eluted by applying buffer A supplemented with 250 mM imidazole. Subsequently, the sample was concentrated and then buffer exchanged using a Nap10 (GE healthcare) column equilibrated in buffer B (20 mM Tris pH 8.0, 150 mM NaCl, 10% glycerol, 2mM DTT).

Over-expression of ΔNp73α and TAp73α proteins (full-length and truncated constructs) fused to MBP was carried out overnight in *E. coli* BL21 DE3 cells at 15 °C. Cell pellets (50 ml expressions) were resuspended in lysis buffer (20 mM Tris pH 8.0, 250 mM NaCl, 10% glycerol, 0.2% NP-40, 2mM DTT, lysozyme, 100 μg/ml DNAse, 100 μg/ml RNAse, cOmplete EDTA-free (Roche)), lysed by sonication and cleared by centrifugation. Supernatants were then incubated with 100 μl of pre-equilibrated amylose resin beads for 2 hours at 4°C. Subsequently, resin was extensively washed with PD buffer (20 mM Tris pH 8.0, 150 mM NaCl, 2mM DTT, cOmplete EDTA-free). For the pulldown experiment, 10 μl of amylose resin coupled to MBP- p73/ΔNp73α proteins were incubated with clarified lysates of HEK293T expressing 3xFlag- E2F4 or recombinant purified E2F4*s*/DP1*s* heterodimer for 2 hours at 4°C. After two quick washing steps with PD buffer, complexes were eluted by incubation with 20 μl of PD buffer supplemented with 20 mM maltose for 15 minutes at 4°C. PD reactions were migrated onto two separate 10% SDS-PAGE gel. One gel was used for Western blot to detect E2F4 or E2F4*s*/DP1*s*, the other for Coomassie staining to detect MBP-p73/ΔNp73α proteins.

#### Immunoblotting

Western blot detection of endogenous proteins was performed using the following antibodies: p73 (Abcam, ref: ab215038), E2F4 (Santa Cruz Biotechnology, ref. sc-398543X), E2F5 (Genetex, ref. GTX129491), DP1 (Abcam, ref. ab124678), p130 (Cell Signaling, ref. 13610S) β-actin (clone C4, MP Biomedicals), GAPDH (6C5, ref. sc-32233, Santa Cruz). Detection of tagged constructs was done using: HA-peroxidase antibody (Roche, ref: 12013819001), anti-TAP antibody (Thermofisher Scientific, ref. CAB1001), anti-Flag antibody (Sigma, ref. F3165) antibody and anti-Gluc antibody (New England Biolabs, ref. E8023).

#### mRNA-seq

38HK cells transfected with scramble siRNA or E2F4-5 siRNAs were collected at 48 hours after transfection. Total RNA was extracted from 38HK (about 10^6^ cells per sample) using the RNeasy Mini kit from QIAGEN and quantified by Qubit.

A total of 6 samples (3 for scramble siRNA and 3 for E2F4-5 siRNA) were analyzed by the GenomEast platform of IGBMC (Illkirch, France). RNA-seq libraries were generated from 500 ng of total RNA using the TruSeq Stranded mRNA Library Prep Kit and TruSeq RNA Single Indexes kits A and B (Illumina, San Diego, CA), according to the manufacturer’s instructions. The read length was 50 nt. The mean total reads per sample was 59,103,999.

Mapping of the reads was processed with STAR 2.7.3a on the primary assembly of the latest release of the human genome (67) (GRCh38.p13, release 33, PRI version: https://www.gencodegenes.org/human/) with corresponding comprehensive gene annotations.

No soft clipping was accepted. Of the reads, 79% mapped once on the genome, 14% multiple times and 6.4% were below the minimum length threshold to map. Reads were counted using htseq-count version 0.11.2 (68) with reverse strand matching (option “stranded reverse”).

Differential expression analysis was done with DESeq2 1.24.0 (69) with Benjamini-Hochberg correction for multiple tests on R 3.6.2.

#### RT-qPCR

Total RNA was extracted from cultured cells using the NucleoSpin RNA II Kit (Macherey-Nagel). The RNA obtained was reverse-transcribed to cDNA using the RevertAid H minus First Strand cDNA Kit (Life Technologies) according to the manufacturer’s protocols. Real-time quantitative PCR (qPCR) was performed using the LightCycler 480 SYBR Green I Master (Roche) or the Mesa Green qPCR MasterMix Plus for SYBR Assay (Eurogentec) with the primers listed in S6 Table. Primers were selected on PrimerBank database (70). Reactions were run in triplicate and expression was normalized to GAPDH. The expression analysis was performed using the MxPro QPCR software (Agilent).

#### Chromatin immunoprecipitation (ChIP)

ChIP was performed using the Shearing ChIP and OneDay ChIP kits (Diagenode) according to the manufacturer’s instructions. Briefly, cells were sonicated to obtain DNA fragments of 200– 500 bp. Sheared chromatin was immunoprecipitated with antibodies against the following proteins/tags: HA (Abcam, ref. ab9110), p73 (Abcam, ref. ab215038), E2F4 (Santa Cruz Biotechnology, ref. SC-398543X), p130 (Cell Signaling, ref. 13610S). 10% of the sheared chromatin was kept as the input for the ChIP.

Immunoprecipitated chromatin has been analysed by q-PCR using the LightCycler 480 SYBR Green I Master (Roche) on a LightCycler® 96 Instrument or the Mesa Green qPCR MasterMix Plus for SYBR Assay (Eurogentec) on a Stratagene Mx3005P Multiplex Quantitative Real Time PCR System. The sequences of primers used for qPCR are described in S6 Table. Primers surrounding the target region were checked for specificity using the NCBI Primer designing tool. ChIP qPCR results were analyzed by evaluating signal of enrichment over noise normalized to Input.

#### Quantification and statistical analysis

Quantification of protein levels from western blot bands was done using the Evolution-Capt Edge software (Vilber) or ImageLab software (Biorad). The data presented are expressed as means ± SD. *P*-values are calculated using unpaired Student’s t-test.

### DATA AVAILABILITY

The mass spectrometry proteomics data have been deposited to the ProteomeXchange Consortium via the PRIDE (71) partner repository with the dataset identifier PXD022947. The RNAseq analyses have been deposited to the GEO database (72) with the identifier GSE162816. Original data files for Western blot analyses are deposited on the public repository

Mendeley Data (doi: 10.17632/cd5hsz8z8w.1).

## AUTHORSHIP CONTRIBUTIONS

**Valerio Taverniti:** Conceptualization, Investigation, Formal analysis, Visualization. **Hanna Krynska:** Investigation, Formal analysis, Visualization. **Assunta Venuti:** Investigation. **Marie-Laure Straub:** Investigation, Methodology. **Cécilia Sirand:** Investigation, Methodology. **Eugenie Lohmann:** Data curation, Formal analysis. **Maria Carmen Romero- Medina:** Investigation. **Stefano Moro:** Investigation. **Alexis Robitaille:** Formal analysis. **Luc Negroni:** Methodology, Formal analysis. **Denise Martinez-Zapien:** Investigation. **Murielle Masson:** Methodology. **Massimo Tommasino:** Conceptualization, Supervision, Manuscript writing, Project administration, Funding acquisition. **Katia Zanier:** Conceptualization, Formal analysis, Visualization, Manuscript writing, Project administration, Funding acquisition.

## ACKNOWLEDGEMENTS

The authors would like to thank Bertrand Seraphin and Christophe Romier (IGBMC, Strasbourg), and Yves Jacob (Institut Pasteur, France) for providing expression vectors. The authors are grateful to Gunter Stier (BZH, University of Heidelberg), Christian Gaiddon (INSERM U1113, Strasbourg), the engineers of the GenomEast platform of IGBMC (Christelle Thibault-Charpentier and Bernard Jost), and members of the IARC and BSC-UMR7242 teams for precious help and advice.

This work received institutional support from IARC/WHO, CNRS, and Université de Strasbourg. The work was supported by grants from ‘Fondation ARC pour la recherche sur le cancer’ (ref. PJA 20151203192) and the ‘Institut National de la Santé et de la Recherche Médicale’ (ref. ENV201610) to MT, and Agence Nationale de la Recherche (ANR JCJC, ref. ANR-13-JSV8-0004-01), ‘Ligue contre le Cancer’ (Région Grand Est - CCIR), ‘Fondation pour La Recherche Medicale’ (Equipes FRM, ref. DEQ20180339231), and ‘Alsace contre le Cancer’ to KZ. This research was partially funded by the Italian Ministry of Health (to MT), Ricerca Corrente 2022, Del. 219/2022.

Where authors are identified as personnel of the International Agency for Research on Cancer/World Health Organization and the Istituto Tumori Giovanni Paolo II, the authors alone are responsible for the views expressed in this article and they do not necessarily represent the decisions, policy, or views of the institutions with which they are affiliated.

## LEGENDS FOR SUPPLEMENTARY FIGURES

**S1 Figure.** Complex immunoprecipitation using an anti-E2F4 antibody. (**A**) IP experiment on total 38HK nuclear extracts using either IgG or an anti-E2F4 antibody. Detection of endogenous ΔNp73α, E2F4 and p130 proteins was done by Western blotting using anti-p73, anti-E2F4 and anti-p130 antibodies, respectively. (**B**) Sucrose gradient/co-IP experiments performed on 38HK nuclear extracts stably expressing HA-ΔNp73α. Indicated fractions were immunoprecipitated using anti-E2F4 antibody and analyzed by Western blot using antibodies recognizing the HA tag, E2F4, and p130 proteins, respectively. Unfractionated nuclear extract (NE) was loaded on the same gel as a reference for protein migration. Images for NE and sucrose fractions are derived from different exposures of the same membrane (see original Western blot image on Mendeley data). Note that the p130 (highlighted by a red Asterix) is partially masked by a closely migrating non-specific band.

**S2 Figure.** Size exclusion chromatography/co-IP experiments. (*Top panel*) 38HK nuclear extracts stably expressing HA-ΔNp73α were applied to a Superose 6 10/300 increase column. (*Middle panel*) Fractions 20 to 30 were found to contain ΔNp73α, E2F4 and p130 proteins. Gel filtration fractions were migrated on a 10% SDS-PAGE gel and analyzed by Western blot using antibodies recognizing the HA tag, E2F4, and p130 proteins. (*Bottom panel*) Fractions 20 to 30 were immunoprecipitated using anti-HA antibody-coupled agarose beads and analyzed for HA-ΔNp73α, E2F4, and p130 proteins.

**S3 Figure.** Sucrose gradient/co-IP experiments in conditions of single E2F4 or E2F5 knockdown. 38HK were transfected with siRNAs against E2F4 (**A**) or against E2F5 (**B**). (*Left panels)* Sucrose fractions. (*Right panels*) Immunoprecipitations of sucrose fractions using an anti-HA antibody. See also legend of Fig. 1D of the main text.

**S4 Figure.** Comparison of ΔNp73α interactions with E2F4 and E2F5. (*Left panel*) GPCA analysis of Gluc1-ΔNp73α *versus* Gluc2-E2F4 and Gluc2-E2F5. (*Right panel)* Expression levels of Gluc1-ΔNp73α, Gluc2-E2F4 and Gluc2-E2F5 in HEK293T cells. Note that the differences in binding responses between E2F5 and E2F4 are related to the different expression levels of the two fusion proteins.

**S5 Figure.** Expression level of E2F4, E2F5, TDP1 and p130 in normal and tumor tissues. The expression matrix plot of E2F4, E2F5, TDP1 and p130 in tumors (T) or normal tissues (N) has been created with the GEPIA2 tool (http://gepia2.cancer-pku.cn)- Multiple Gene Analysis-Multiple Gene Comparison. Gene list: E2F4, E2F5, TDP1 and p130. Dataset: all cancer types. Matched Normal data: TCGA tumor + TCGA normal + GTEx normal. The density of color in each block represents the median expression value of the indicated gene in a given tissue, normalized by the maximum median expression value across all blocks. Different genes in same tumors or normal tissues are compared. GTE: Genotype-Tissue Expression. **ACC**: Adrenocortical carcinoma, **BLCA**: Bladder Urothelial Carcinoma, **BRCA**: Breast invasive carcinoma, **CESC**: Cervical squamous cell carcinoma and endocervical adenocarcinoma, **CHOL**: Cholangio carcinoma, **COAD**: Colon adenocarcinoma, **DLBC**: Lymphoid Neoplasm Diffuse Large B-cell Lymphoma, **ESCA**: Esophageal carcinoma, **GBM**: Glioblastoma multiforme, **HNSC:** Head and Neck squamous cell carcinoma, **KICH**: Kidney Chromophobe, **KIRC**: Kidney renal clear cell carcinoma, **KIRP**: Kidney renal papillary cell carcinoma, **LAML**: Acute Myeloid Leukemia, **LGG**: Brain Lower Grade Glioma, **LIHC**: Liver hepatocellular carcinoma, **LUAD**: Lung adenocarcinoma, **LUSC**: Lung squamous cell carcinoma, **OV**: Ovarian serous cystadenocarcinoma, **PAAD**: Pancreatic adenocarcinoma, **PCPG**: Pheochromocytoma and Paraganglioma, **PRAD**: Prostate adenocarcinoma, **READ**: Rectum adenocarcinoma, **SARC** : Sarcoma, **SKCM**: Skin Cutaneous Melanoma, **STAD**: Stomach adenocarcinoma, **TGCT**: Testicular Germ Cell Tumors, **THCA**: Thyroid carcinoma, **THYM**: Thymoma, **UCEC**: Uterine Corpus Endometrial Carcinoma, **UCS**: Uterine Carcinosarcoma.

**S6 Figure.** E2F4-5 knockdown in primary HPKs. E2F4-5 levels in HPK cultures treated with SC (white histograms) or siE2F4-5 (black histograms) RNAs. Total proteins used for the Western blot have been extracted using the Nucleospin RNA/Protein kit (Macherey-Nagel) following the manufacturer’s protocol. Histograms show averages derived from three independent experiments, with error bars representing standard deviations. Protein levels are normalized to actin and then to the control SC condition.

**S7 Figure.** SAHF formation upon E2F4 or E2F5 silencing by shRNA in 38HK. (**A**) mRNA levels of E2F4 and E2F5 genes in conditions of knockdown using PLKO.1 plasmids containing the shE2F4 or shE2F5 sequences (i.e. shE2F4 and shE2F5) as measured by RT-qPCR. The pLKO.1 empty plasmid was used as negative control (*pLKO.1*). (**B**) (*Upper panel*) Representative immunofluorescence images of 38HK cells transfected with *pLKO.1*, shE2F4 or shE2F5. At 48 hours post-transfection, cells were layered on coverslips coated with polylysine and probed using an anti-H3K9me3 antibody followed by secondary Alexa Fluor 488-conjugated antibody. Nuclei were stained with DAPI (coloured blue). Images were merged using ImageJ software. (*Lower panel*) Quantification of cells positive for H3K9me^3^ marks in the four conditions was performed using ImageJ software. > 100 cells considered for each sample. (**A**-**B**) The data are averages of three independent experiments. *P*-values are obtained from unpaired *t*-test, *n* = 3 biological triplicates (**: *P* < 0.01; ***: *P* < 0.001).

**S8 Figure.** Ectopic expressions of E2F4/5 decreases SAHF formation in 38HK transfected with shE2F4/5 plasmids. (*Upper panel*) Representative immunofluorescence images of 38HK cells transfected with different combinations of *pLKO.1* or *pCNA* empty control plasmids, pLKO.1 shE2F4 or shE2F5 plasmids (i.e. shE2F4 or shE2F5), and pCNA plasmids containing the cDNAs of E2F4 or E2F5 (i.e. E2F4 and E2F5). For each condition two plasmids are transfected simultaneously. At 48 hours post-transfection, cells were layered on coverslips coated with polylysine and probed using the anti-H3K9me3 and Alexa Fluor 488-conjugated antibodies. (*Lower panel*) Quantification of cells positive for H3K9me^3^ marks in the different conditions. > 100 cells considered for each sample. The data are averages of three independent experiments. *P*-values are obtained from unpaired *t*-test, *n* = 3 biological triplicates (*: *P* < 0.05; ***: *P* < 0.001).

**S9 Figure.** E2F4-5 and ΔNp73α knockdown in 38HK. **(A)** Western blot analysis of E2F4 and E2F5 gene expression in the 38HK cultures treated with SC (white histograms, *lower panel*) or siE2F4-5 RNAs (black histograms, *lower panel*) and used for mRNA-seq studies (Fig. 6A and B of the main text). All samples were run on the same gel (see also Mendeley for original image). (**B**) Western blot analysis of ΔNp73α, E2F4 and p130 protein levels in 38HK cultures treated with sense (S, white histograms, *lower panel*) or antisense (AS, black histograms, *lower panel*) oligonucleotides targeting ΔNp73α. The three 38HK cultures were used for the gene expression analysis shown in Fig. 6C of the main text. Total proteins were extracted using the Nucleospin RNA/Protein kit (Macherey-Nagel). (**A-B**) Histograms report averages that are normalized to the respective SC and S control conditions, with error bars representing standard deviation values from three independent experiments. Protein levels are additionally normalized to GADPH.

**S10 Figure.** E2F4-5 and ΔNp73α knockdown in cancer cells. **(A)** RT-qPCR analysis of E2F4 and E2F5 expression in HNC-136 and CAL-51 cell cultures treated with SC (white histograms) or siE2F4-5 (black histograms) siRNAs and used for the gene expression analysis shown in Fig. 8 of the main text and S8 Fig. (**B**) Evaluation of cell death in CAL51 cultures treated with S and AS against ΔNp73α according to the optimized protocol. Total cells (adherent and floating) were colored with trypan blue. **(C)** Western blot analysis of ΔNp73α protein levels in CAL-51 cultures treated with sense (S) or antisense (AS) oligonucleotides against ΔNp73α. These CAL- 51 cultures were used for the gene expression analysis shown in Fig. 8 of the main text. Total proteins were extracted using the Nucleospin RNA/Protein kit (Macherey-Nagel). (**A** and **C**) Histograms report averages are normalized to the respective SC and S conditions, with error bars representing standard deviations from three independent experiments. Protein levels are additionally normalized to actin.

**S11 Figure.** Expression profiles of selected genes in HNC-136 cells. mRNA levels of selected genes determined by RT-qPCR in conditions of E2F4-5 knockdown. The red bar corresponds to the expression level of each gene in the scramble or sense control conditions (set to 1). “E2F4-5 siRNA/scramble” values > 1 and < 1 indicate upregulation and downregulation of gene expression, respectively. Error bars report on the standard deviations from three independent experiments. *: *P* < 0.05; **: *P* < 0.01. See also legend of Fig. 6 of the main text.

## LEGENDS FOR SUPPLEMENTARY TABLES

**S1 Table.** Nuclear protein binding partners of ΔNp73α. List of proteins displaying an enrichment of at least 10-fold in the ΔNp73α-TAP proteomics experiment compared with control (TAP-only) conditions. All proteins listed have been detected by more than two unique peptides, with the exception of DP1 that was detected by two unique peptides.

**S2 Table**. mRNA-seq analysis: genes significantly upregulated upon E2F4-5 knockdown in 38HK. Significantly upregulated genes are associated with adjusted *P*-values ≤ 0.05. Genes are ranked based on adjusted *P*-values. The presence or absence of E2F4 and TP53/TP73 RE within the long promoter region (+2500 nt from TSS) is reported for each gene.

**S3 Table.** mRNA-seq analysis: genes significantly downregulated upon E2F4-5 knockdown in 38HK. Significantly downregulated genes are associated with adjusted *P*-values ≤ 0.05. See also legend of S2 Table.

**S4 Table.** Results of E2F4 and p53/p73 RE predictions for genes upregulated upon E2F4-5 knockdown in 38HK. Sequences of upregulated genes were recovered from the Ensembl database version 86 dataset and biomaRt 2.42.0. The 2500 nt region upstream of TSS (defined as the long promoter region) was searched for E2F4 and TP53/TP73 REs using the JASPAR2020 (76) (https://github.com/da-bar/JASPAR2020) 0.99.8 and TFBSTools 1.25.2 packages. The score threshold was set to 0.85.

**S5 Table.** Effects of E2F4-5 and ΔNp73α knockdown on the expression of selected genes in 38HK. Genes which are significantly upregulated in the RNA-seq analysis and that have been tested in classical RT-qPCR experiments. +: rescue associated with significant *p-*value; -/+: rescue associated with higher average mRNA levels but not significant *p*-values; -: no rescue.

**S6 Table.** Oligonucleotide sequences for gene knockdown, RT-qPCR, and ChIP experiments.

